# Structural Basis for pre-tRNA Recognition and Processing by the Human tRNA Splicing Endonuclease Complex

**DOI:** 10.1101/2022.09.02.506201

**Authors:** Cassandra K. Hayne, Kevin John U. Butay, Zachary D. Stewart, Juno M. Krahn, Lalith Perera, Jason G. Williams, Robert M. Petrovitch, Leesa J. Deterding, A. Gregory Matera, Mario J. Borgnia, Robin E. Stanley

## Abstract

Across all walks of life, certain transfer RNA (tRNA) transcripts contain introns. Pre-tRNAs with introns require splicing to form the mature anticodon stem loop (ASL). In eukaryotes, tRNA splicing is initiated by the heterotetrameric tRNA splicing endonuclease (TSEN) complex. All TSEN subunits are essential and mutations within the complex are associated with a family of neurodevelopmental disorders known as pontocerebellar hypoplasia (PCH). The pathogenesis of PCH is poorly understood. Moreover, a lack of structures for any eukaryotic TSEN complex has hindered our understanding of tRNA recognition and processing. Here, we report Cryo-Electron Microscopy (cryo-EM) structures of the human TSEN•pre-tRNA complex, trapped in the pre-cleavage state, at near atomic resolution. These structures reveal the overall architecture of the complex, along with extensive tRNA binding interfaces within the complex. Although it shares structural homology with archaeal TSENs, the human TSEN complex contains additional features important for recognizing the acceptor stem and D-arm of the pre-tRNA. Our findings also establish the TSEN54 subunit as more than a simple molecular ruler; it functions as a pivotal scaffold for the pre-tRNA and the two endonuclease subunits, TSEN2 and TSEN34. Finally, the human TSEN structures enable detailed visualization of the molecular environments of PCH-causing missense mutations, providing crucial insight into the mechanism of eukaryotic pre-tRNA splicing and neurodevelopmental disease.

## Main

Prior to their involvement in translation, tRNAs undergo extensive processing and modification^1^. Although most tRNA genes are intronless, a subset of pre-tRNAs contain an intron that forms base pairing interactions with the anticodon^2^. These intervening sequences must be removed from the precursor prior to translation. Within the human genome, there are several isodecoder families that all contain introns; thus tRNA splicing is essential^3,4^. Eukaryotic tRNA intron cleavage is catalyzed by the heterotetrameric TSEN complex^5^. The TSEN complex contains two presumptive structural subunits, TSEN54 and TSEN15, along with two endonuclease subunits, TSEN34 and TSEN2, that catalyze cleavage at the 3’ and 5’-splice sites, respectively^6,7^. Beyond providing structural support, the TSEN54 subunit is thought to function as a molecular ruler that properly positions the 5’-splice site for cleavage by TSEN2^8-10^. However, the overall molecular architecture of the TSEN complexes of both yeast^6^ and humans^7^ has remained elusive. The only available TSEN structures include an NMR structure of TSEN15^11^, and a crystal structure of TSEN15 bound to the C-terminal region of TSEN34^12^. The eukaryotic TSEN proteins are thought to have evolved from an archaeal ancestor called EndA, which forms homo-oligomers that recognize a stereotypical bulge-helix-bulge (BHB) RNA motif. How the eukaryotic TSEN complexes evolved to accommodate recognition of full pre-tRNA substrates is unknown^7,13-15^.

The human TSEN complex is known to associate with the polyribonucleotide 5’-hydroxyl-kinase CLP1^7^, which also functions in mRNA 3’-end processing^16^. CLP1’s association with the TSEN complex is perplexing, given that its kinase activity would block RTCB mediated ligation of the exon halves. *In vitro* reconstitution of with the human TSEN complex has established that CLP1 is not required for intron removal^12,17^. *In vivo*, CLP1 likely plays an important regulatory role in tRNA splicing by inhibiting exon ligation and/or targeting exonucleolytic decay of the intron^2,5,17^. Mutations in CLP1 and all four TSEN subunits are associated with multiple subtypes of the PCH family of inherited neurodevelopmental disorders^18-21^. PCH subtypes are characterized by genotype and clinical features including microcephaly, intellectual disability, locomotor dysfunction and premature death; however, the etiology of these diseases remains poorly understood^22^.

Here, we report near atomic resolution cryo-EM structures of the human TSEN complex bound to intron-containing pre-tRNAs. The structures unveil the arrangement and function of all four subunits and reveal a structural core within the complex that is conserved from archaeal ancestors. The structures also identify extensive eukaryotic specific interfaces within the complex that mediate recognition of the full-length pre-tRNA.

### Structure of the human TSEN complex

We solved two cryo-EM structures of the human TSEN complex bound to intron-containing pre-tRNA-ARG^TCT^-2-1 (Fig. 1). Two different approaches were used to trap the complex in a pre-cleavage state with the intron intact. First, we purified the recombinant wildtype(wt)-TSEN complex using a multi-cistronic bacterial expression system, which we previously reported is sufficient for intron cleavage^17^. We incubated wt-TSEN with pre-tRNA-ARG, containing a 15-nt intron wherein the 2’-hydroxyl at both the 3’ and 5’ cleavage sites was modified to fluorine to prevent catalysis^23^. The structure of wt-TSEN was solved to a resolution of 3.9 Å (Fig. 1a, Extended Data Table 1) and was sufficient for rigid-body docking of the tRNA and the four TSEN subunits using a combination of previously determined structures and AlphaFold models (Extended Data Fig. 1)^11,12,24,25^. The overall density of the pre-tRNA was well resolved, except in the area surrounding the 5’ cleavage site.

**Fig. 1.**
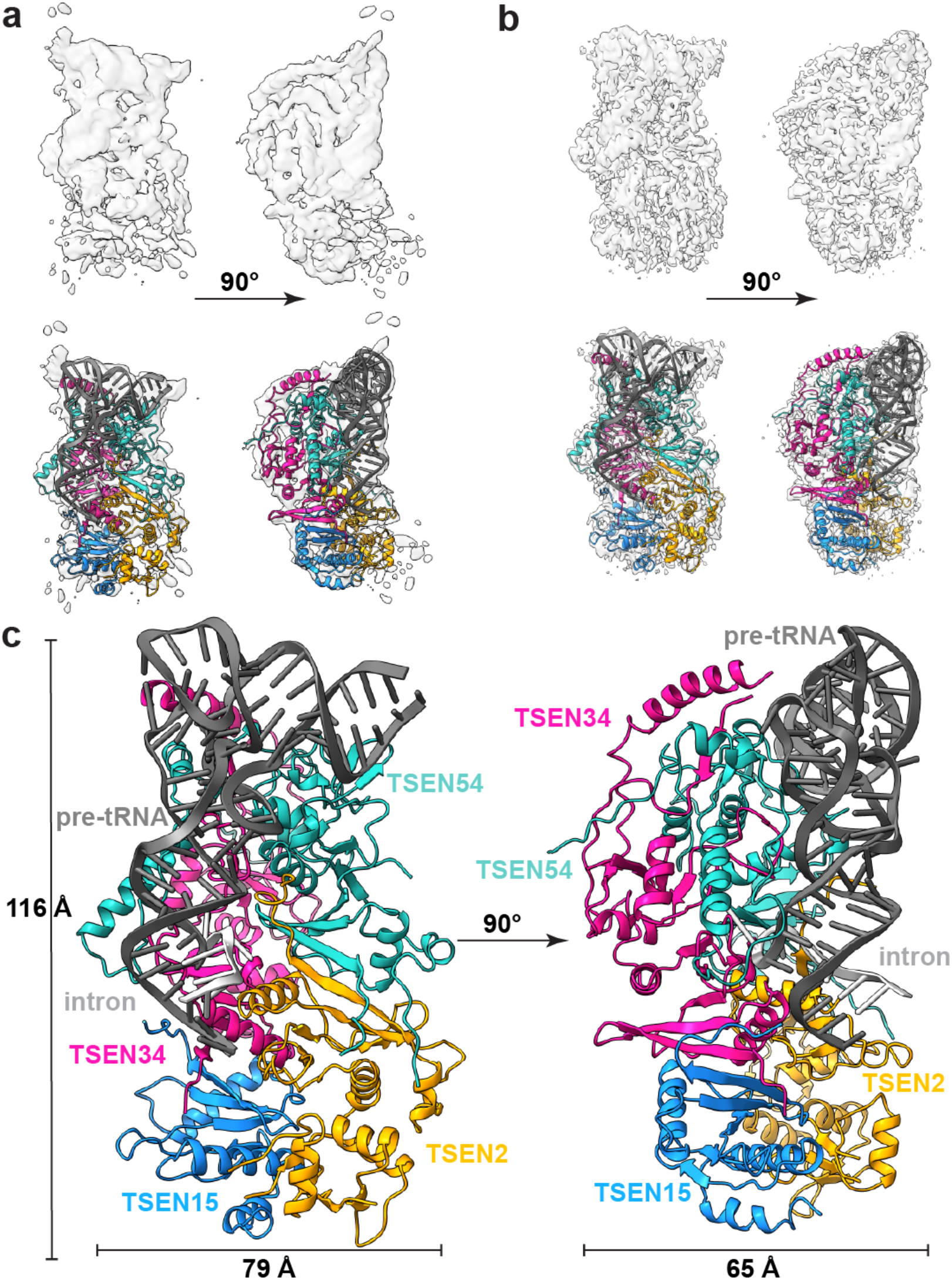
Cryo-EM Structures of the Human TSEN complex. Cryo-EM reconstructions of: **a**. wt-TSEN•pre-tRNA and **b**. endoX-TSEN•pre-tRNA, with cartoon overlays below. Due to its improved resolution, the endoX reconstruction was used to build and refine the overall structure. **c**. Model of the human TSEN complex at 0° and 90°, with overall dimensions of 116 × 79 × 65 Å. The subunits are colored as follows: TSEN54 (teal), TSEN34 (pink), TSEN2 (orange), TSEN15 (blue). The pre-tRNA is shaded in dark grey, with the intron in light grey.

We also solved a structure of the TSEN complex in the presence of the CLP1 RNA kinase. Although the TSEN complex is sufficient for cleaving tRNA introns *in vitro*^12,17^, CLP1 is a well-established interaction partner^16,21^. To prevent intron cleavage, we mutated the three catalytic residues of both TSEN endonucleases. We purified an endonuclease-deficient (endoX)-TSEN•CLP1 complex from a mammalian overexpression system using an affinity tag on CLP1, and incubated the complex with an *in vitro* transcribed unmodified pre-tRNA-ARG ^TCT^-2-1. The structure of the endoX-TSEN•CLP1 complex was solved to an overall resolution of 3.3 Å (Fig. 1b, Extended Data Fig. 2 and Extended Data Table 1). Well resolved density was visible for all four TSEN subunits and the pre-tRNA, but we did not observe any well-ordered density for which we could accurately assign to CLP1. Overall, both the wt-TSEN and endoX-TSEN cryo-EM reconstructions are very similar, and neither contain well resolved density for the intron surrounding the 5’ cleavage site. Due to its improved resolution, the endoX-TSEN•CLP1 reconstruction was used to build the atomic model of the TSEN complex (Fig. 1c, Extended Data Fig. 2).

The four subunits of the TSEN complex form an interconnected, rectangular shaped structure. Overall, the core of the human TSEN complex is structurally similar to the archaeal complexes (Extended Data Fig. 3 a). Previous structural studies of the archaeal enzymes established that the interfaces within the complex are mediated by two key features: a C-terminal β-sheet (β9) and loop (L10)^26^. Both structural features are conserved in the human TSEN complex, reinforcing the hypothesis that the eukaryotic TSEN complex evolved from a common archaeal ancestor^27^. Both the TSEN54-TSEN2 and TSEN34-TSEN15 interactions are mediated by anti-parallel β-β interfaces formed by the final β-strand from each subunit (Extended Data Fig. 3b-d). The same TSEN34-TSEN15 interface was also recently observed in a crystal structure of a portion of the C-terminal domain of TSEN34 bound to TSEN15^12^. The TSEN34-TSEN15 and TSEN54-TSEN2 dimers are held together by the L10 loop interface (Extended Data Fig. 3 b,e,f). The negatively charged L10 loop in TSEN15 (residues 144-150) inserts into a positively charged cleft in the TSEN2 subunit (Extended Data Fig. 3e). Similarly, the L10 loop in TSEN54 (residues 506-509) inserts into a cleft in the TSEN34 subunit (Extended Data Fig. 3f). Beyond these conserved interfaces, the human TSEN complex also contains additional interfaces not found in the archaeal enzymes (Extended Data Fig. 3g). The N-terminus of TSEN34 and TSEN54 contain large insertions with no structural homology to the archaeal enzymes and TSEN54 forms an extensive interface with TSEN34. The TSEN34 and TSEN54 subunits also have the largest buried surface area within the complex. Additionally, while there are no interfaces between TSEN54 and TSEN15 in the structure, TSEN2 and TSEN34 form interfaces near the active sites.

### TSEN54 Mediates CLP1 Binding

We performed cross-linking mass spectrometry analysis with a lysine reactive cross-linker on the recombinant TSEN complex to gain additional insight into the subunit interfaces (Extended Data Fig. 4). Three of the four TSEN subunits contain large, disordered regions that are not visible in the reconstructions^5^. Further, we did not observe well-ordered density for the CLP1 kinase in the reconstruction, suggesting that it either associates with the complex through a dynamic/disordered region, or it dissociates from the complex during cryo-EM grid preparation. We detected several high-confidence crosslinks within regions of the TSEN core that are visible in the structure (Extended Data Fig. 4a). We also detected crosslinks across the disordered regions of TSEN2, TSEN34, and TSEN54 offering additional insight into interactions mediated by these disordered regions (Extended Data Fig. 4a). Crosslinking was carried out in the absence of tRNA, revealing that the core complex likely does not undergo large scale conformational changes upon binding tRNA. This was supported by molecular dynamics (MD) simulations of the complex, with and without RNA bound, in which we did not observe large conformational changes (Extended Data Fig. 5). MD simulations further revealed the occasional loss of the TSEN15 subunit from the rest of the TSEN complex, likely because of its overall negative charge. The RMSD plots for the TSEN complex with and without RNA as well as the RNA bound complex without TSEN15 were stable over the trajectory used for MD simulations and across three replicates (Extended Data Fig. 5a).

We also conducted cross-linking mass spectrometry analysis of the TSEN•CLP1 complex. We observed fewer crosslinks because the overall protein concentration was lower for the TSEN•CLP1 sample, but we detected high confidence crosslinks between CLP1 and both TSEN2 and TSEN54 (Extended Data Fig. 4b-h), suggesting that TSEN2 and TSEN54 may mediate CLP1 binding to the TSEN complex. To test this hypothesis, we carried out immunoprecipitations from cells transfected with CLP1 and either individual TSEN subunits or the full TSEN complex and found that TSEN54 was sufficient to mediate co-immunoprecipitating CLP1 (Extended Data Fig. 6).

### Structure of pre-tRNA reveals similarities to mature tRNAs

Beyond the architecture of the TSEN complex, our cryo-EM reconstruction also revealed the overall structure of an intron-containing pre-tRNA. Mature tRNAs contain several distinct structural features including, the accepter stem, D and TΨC arms, a variable loop, and ASL (Extended Data Fig. 7a). Globally, the structure of TSEN bound pre-tRNA is very similar to the structure of mature tRNA, except for the ASL containing the intron (Extended Data Fig. 7b). Bending in the ASL region of the bound pre-tRNA suggests that the TSEN complex causes a slight melting of the ASL to position it for cleavage. Previous studies have established that eukaryotic introns are located between nucleotides 37 and 38 of the mature tRNA, forming a specific RNA structure known as the bulge-helix-bulge (BHB) or a more relaxed bulge-helix-loop (BHL) motif^2^. The anticodon-intron (A-I) helix contains the anticodon which base pairs with the intron and is flanked by single-stranded bulges/loops where cleavage occurs. Within our cryo-EM reconstructions, the density was sufficient to build a model for the A-I helix along with the bulge around the 3′-cut site (Extended Data Fig. 7c). The loop encompassing the 5′-cut site is unstructured in both cryo-EM reconstructions, suggesting that this portion of the intron is highly flexible. This is not surprising given that the TSEN complex can accommodate large insertions in this loop that do not disrupt cleavage^2,28^.

### pre-tRNA Recognition by the TSEN complex

How the eukaryotic TSEN complex recognizes pre-tRNAs has been a long-standing question. A crystal structure of an archaeal EndA engaging a BHB substrate established how conserved features surrounding the EndA active sites position the bulged residues for cleavage^8^. For EndA enzymes, the BHB motif is strictly required for recognition and cleavage, which is important because archaeal tRNA genes have introns located in various regions of the pre-tRNA^29^. In contrast, the human TSEN complex relies on recognition of additional tRNA features that mediate pre-tRNA recognition and it can cleave tRNA substrates with a more relaxed bulge-helix-loop (BHL) motif^9,30^. However, the eukaryotic complex can also cleave the universal BHB motif independent of the rest of the tRNA suggesting that recognition and cleavage of the BHB motif is a universal feature of TSEN nucleases^31,32^.

The cryo-EM structure revealed an extensive interface formed between the TSEN complex and several regions of the tRNA (Fig. 2a). The TSEN complex primarily interacts with the upside-down L-shaped tRNA along the under-side of the L, at the intersection of the acceptor-stem and stacking of the D-arm and ASL. TSEN54 was previously designated as a ‘molecular ruler’ based on studies demonstrating that insertions in specific regions of the pre-tRNA can impact the cleavage site^8-10^. TSEN54 contains the largest RNA interface and specifically interacts with the acceptor stem and D-arm of the tRNA, predominantly through interactions with the phosphate backbone (Fig. 2b-e). The acceptor stem is positioned along the top of the complex by TSEN54 and there are no interactions with the T-arm. Previous work revealed that while the acceptor stem is not required for intron cleavage, its removal significantly impacts the kinetic parameters of cleavage^33^, confirming the significance of the TSEN54-acceptor stem interface in anchoring the tRNA within the TSEN complex.

**Fig. 2.**
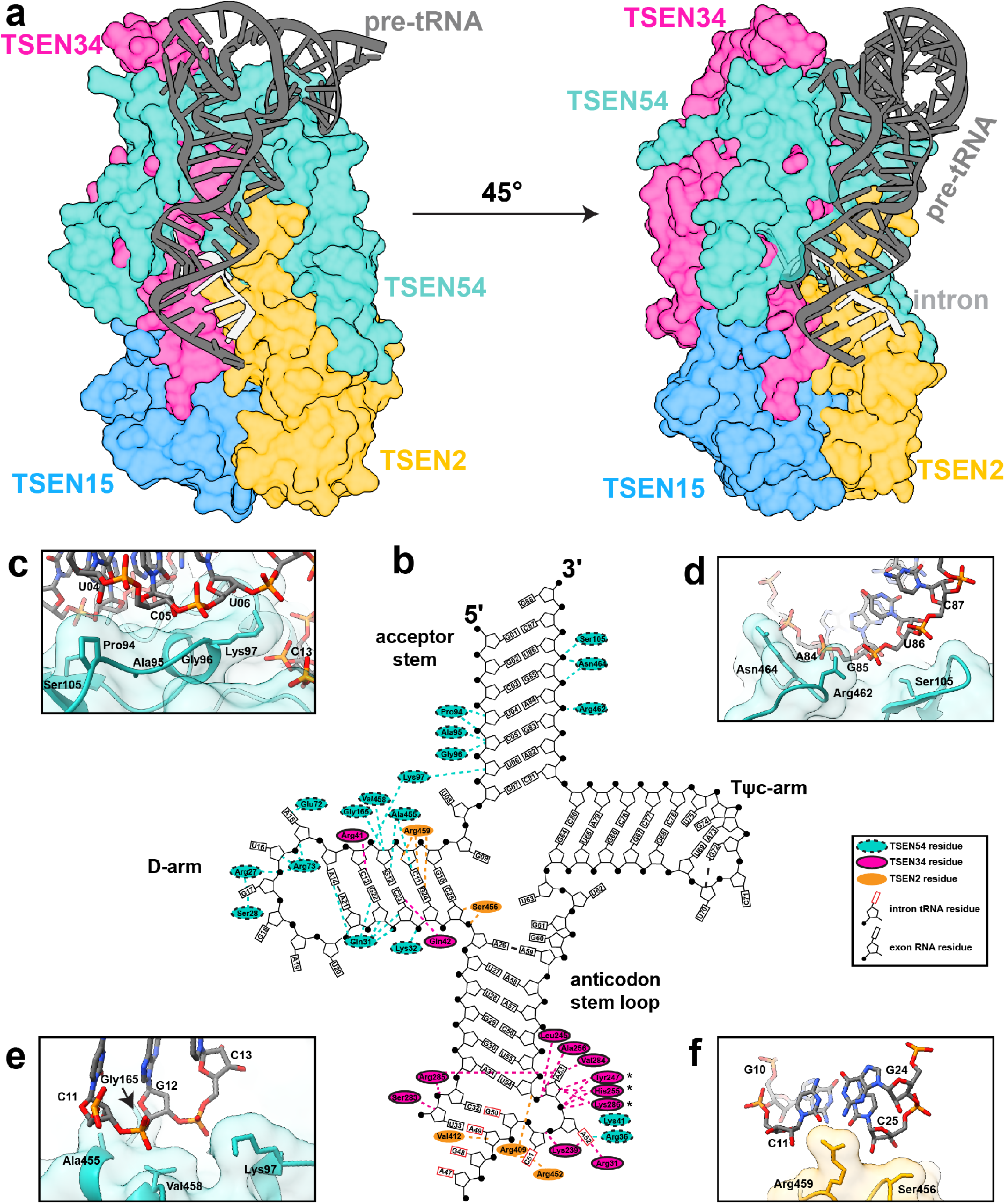
Recognition of pre-tRNA features by the TSEN complex. **a**. Surface representation of the TSEN complex with a cartoon of the pre-tRNA, showing significant interaction between TSEN54 and multiple regions of the pre-tRNA. Colors: Exons (dark grey), intron (light grey), TSEN54 (teal), TSEN34 (pink), TSEN2 (orange), TSEN15 (blue). **b**. Cloverleaf 2D tRNA structure with the TSEN interacting residues labeled. Regions of the tRNA from the intron are indicated with the rectangle, representing the base, boxed in red. Interfaces between TSEN54 and **c**. and **d**. the acceptor stem **e**. the D-arm. **f**. Interface between the D-arm and TSEN2.

Our structure supports previous biochemical data suggesting a central role for TSEN54 in tRNA recognition, but also provides detailed insight into how TSEN54 engages tRNA and supports the overall architecture of the complex. Many of the TSEN54-tRNA interactions are mediated by the N-terminal region of TSEN54, which has no sequence or structural homology to the archaeal EndA proteins (Extended Data Fig. 3g). In addition to TSEN54, the two endonucleases also mediate TSEN-RNA interfaces within the D-arm and the ASL (Fig. 2b,f). Similar to TSEN54, TSEN34 contains several insertions, not found in archaeal EndA, that mediate tRNA binding. These include a loop (TSEN34 residues 33-48) in the N-terminal domain that interacts with the D-arm, and a large alpha-helix positioned near the T-arm. The local resolution of this alpha-helix is poor, suggesting it is dynamic.

To further highlight the significance of the RNA-protein interfaces within the cryo-EM structure, we estimated the free energy of binding by the Molecular Mechanics-Generalized Born and Surface Area (MM-GBSA) approach using the configurations collected from our MD simulations. The calculated free energy of binding for residues from TSEN2, TSEN34, and TSEN54 is shown in Extended Data Table 2. From this analysis, we observed that TSEN54 provides the largest contribution to RNA binding. Collectively, our data suggest that TSEN54, TSEN34, and TSEN2 are each important for recognition of the tRNA. Although not visible in the structure, we hypothesize that TSEN15 likely mediates interactions with the intron surrounding the 5′ splice site.

### The TSEN complex has a conserved catalytic core

The cryo-EM structure also provided insight into cleavage at the 3′ splice site (Fig. 3a). The 3-nt bulge in the 3′ splice site is well ordered in our cryo-EM structure, in contrast to the 5′ splice site. Analogous to the archaeal EndA•BHB structure, the 3-nt bulge forms an open knot-like conformation, presumably to orient the RNA for cleavage^8^. This interaction is mediated primarily by residues from TSEN34 including V284 and K239, but is further mediated by TSEN2(R409, R452) and TSEN54(R36, K41) (Fig. 3b). The close proximity of residues from TSEN54 to the 3′ splice site appears to be a feature unique to the eukaryotic TSEN complex. To prevent cleavage, we mutated the three catalytic residues from TSEN34 (H255, K286, Y247) to alanine, however in our reconstruction we observe a low occupancy of these active site residues, sufficient to roughly model side chain positions for these residues (Fig 3c). This heterogeneity is the result of purifying the TSEN complex, using CLP1 as bait, from a mammalian expression line containing the endogenous enzymes. We attempted to resolve this heterogeneity through cryo-EM data processing but were unsuccessful at generating a high-resolution map with strong density for the WT TSEN34 catalytic residues, most likely because the population of these particles is very small. Superposition of the EndA•BHB structure (Extended Data Fig. 8a) reveals that these catalytic residues are in similar positions (Extended Data Fig. 8b-e).

**Fig. 3.**
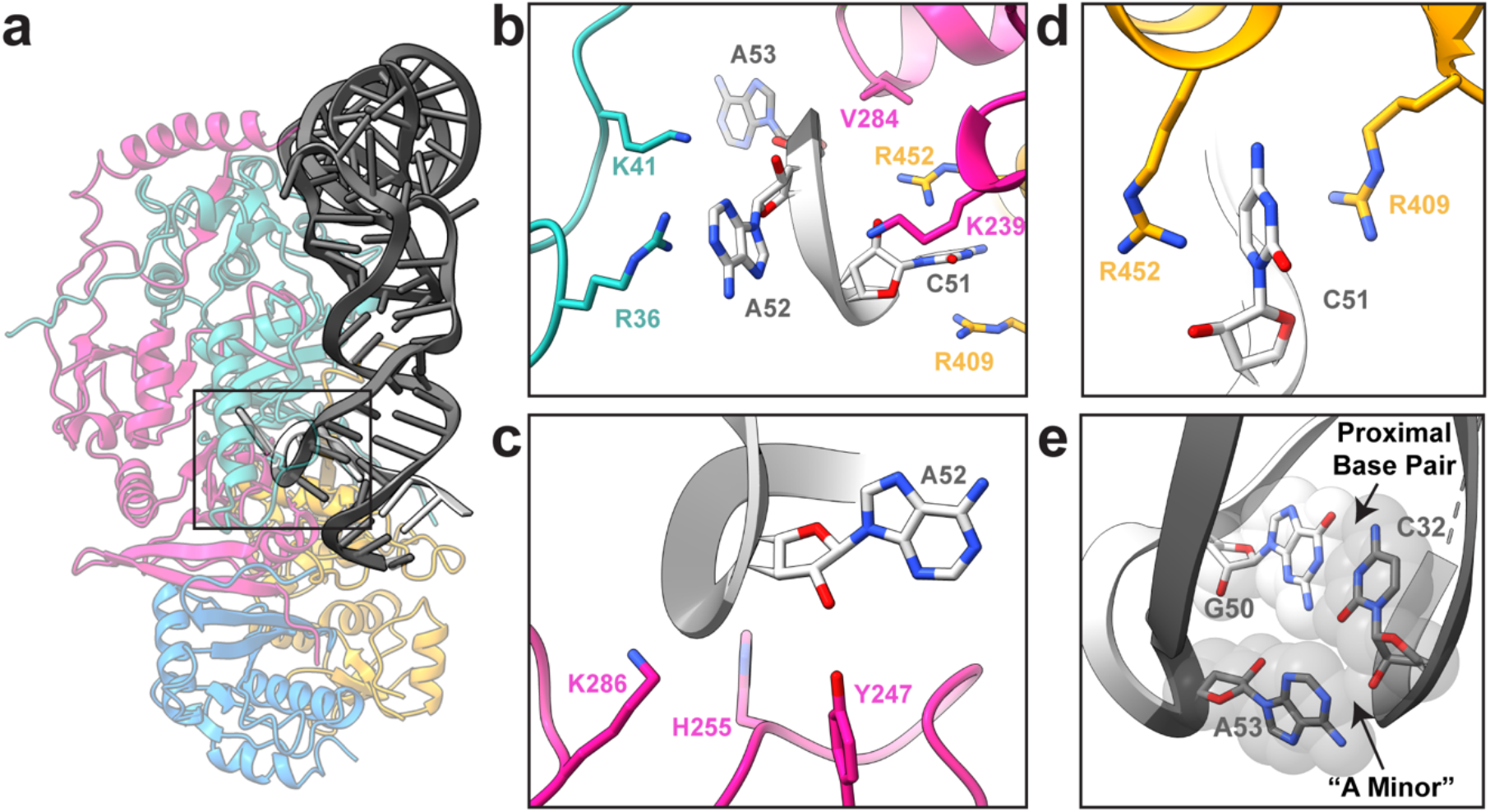
Conserved catalytic core at the 3’ splice site. **a**. Cartoon structure of the complex with a box highlighting the 3’ splice site. **b**. Overview of the three nucleotide bulge in the 3’ splice site with important interactions labeled including TSEN54 (teal) (R36 and K41), TSEN34 (pink) (V284 and K239), and TSEN2 (orange) (R409 and R452). **c**. Modeled positions of the catalytic residues from TSEN34 (Y247, H255, and K286), which were mutated to alanine to prevent cleavage. **d**. TSEN2 residues R409 and R452 form a cation-π interaction with the C51 RNA base of the intron to stabilize the bulge. **e**. The ASL contains the proximal base pair (C32 and G50), important for splicing *in vivo*. The C32 base forms an A-minor like motif with A53 from the bulge.

The 3′ splice site is further supported by a cross-subunit interaction that stabilizes the first bulged nucleotide. The EndA•BHB crystal structure revealed that there is cooperativity between the splice sites in archaeal nucleases that is mediated by a cross-subunit cation-π interface^8^. The bulged nucleotide is held in place by a pair of residues (R:R or R:Y/W) from the opposite endonuclease, forming a cation-π sandwich. Earlier studies of the yeast TSEN complex suggested that this cross-subunit cooperativity was required for cleavage at the 5′ splice site but dispensable for cleavage at the 3′ splice site^34^. Sequence comparisons across multiple EndA/TSEN nucleases suggest that the equivalent residues in human TSEN2 that form the cation-π interface are R409 and W453^34^; however, our structure revealed that R452, not W453, mediates the cation-π sandwich in the 3′ splice site (Fig. 3d, Extended Data Fig. 8c). Although we do not observe tRNA intron density in the 5′ splice site, we modeled RNA into the active site by superposition with the archaeal EndA BHB structure. We observe that the TSEN34 cation-π interface is formed by TSEN34 residues R279 and W306 (Extended Data Fig. 8d) and this superposition enabled us to roughly model the TSEN2 catalytic residues (Extended Data Fig. 8e). Both the EM-structure and modeling of the TSEN2 active site suggests that cross-subunit stabilization is a feature conserved across EndA/TSEN endonucleases.

We determined the significance of the human TSEN34 and TSEN2 cation-π interfaces in mediating tRNA cleavage within the human complex using nuclease assays. We generated single and double mutants of the TSEN2 (R409A, R452A) and TSEN34 (R279A, W306A) and isolated the TSEN complex containing the specific mutants by co-IP with an affinity tag on either TSEN2 or TSEN34 (Extended Data Fig. 8 f,g). We incubated resin bound TSEN complex with a pre-tRNA containing the broccoli aptamer in the intron and then ran the quenched reaction on a denaturing gel. All of the single mutants were active, but the TSEN34 (R279A/W306A) double mutant abolished cleavage at the 5′ splice site (Extended Data Fig. 8g), supporting earlier work with the yeast complex^34^. In contrast, the TSEN2 (R409A/R452A) mutant retained activity (Extended Data Fig. 8f), revealing that this cation-π interface is dispensable for cleavage at the 3′ splice site^34^. Whereas the TSEN2 cation-π interface at the 3′ splice site is not required for pre-tRNA cleavage, its conservation suggests it plays an important role. We hypothesize that the TSEN2 cation-π interface is not essential because of the extensive contacts that can hold the pre-tRNA in the absence of the cation-π sandwich, although this interface may support cleavage of non-tRNA substrates. Given that the 5′ splice site is not well ordered in our structure, it likely has a weaker RNA interface and requires the cation-π sandwich.

The TSEN structure further revealed that the 3′ bulge is supported by the well characterized anticodon-intron interaction known as the proximal base-pair (PBP, C32-G50). Previous work has established the significance of the PBP in mediating cleavage at the 3′ splice site^28,35^. In Drosophila and human intronic tRNAs, this base pair is strictly a C-G, and the strength of the PBP impacts cleavage^4,5,28^. Analogous to the EndA•BHB structure, we observe that the PBP forms an A-minor interaction with the last nucleotide in the 3′ bulge, A53 (Fig. 3e). This A-minor motif stabilizes A53 within the 3′ splice site. Within the EndA•BHB structure, this motif is further supported by base stacking with the catalytic histidine^8^. Although the corresponding residue in TSEN34 (H255) is mutated in our structure, given its equivalent position to the archaeal enzymes, we assume it plays a similar role (Extended Data Fig. 8b). Overall, the structure reveals a remarkable amount of similarity between the 3′ splice sites of TSEN/EndA nucleases, supporting earlier work establishing that these enzymes employ common mechanisms to recognize the universal BHB motif and cleave the RNA by an RNAse-A like transesterification reaction^31^.

### Structure reveals location of many PCH disease mutations

We mapped known PCH disease-associated mutations onto our TSEN structure, with the exception of TSEN54(A307S) which lies in a disordered region of TSEN54 not in the structure (Figure 4A)^5^. Interestingly, none of the mutations appear within active site regions of TSEN34 or TSEN2, but instead are found on surface exposed interfaces that may be sites of interaction with other proteins or in regions that appear important for complex stability. For example, we predict that TSEN54-Y119 will decrease protein stability as Y119 is buried within the core of the TSEN54 N-terminal domain. TSEN54-Y119 could also potentially interfere with tRNA binding as it lies within hydrogen bonding distance of TSEN34-R41 which associates with the D-arm of the pre-tRNA. We predict that TSEN34(R58W) would decrease stability with the TSEN34 subunit as this would disrupt the salt bridge formed between TSEN34-R58 and TSEN34-E218. This supposition agrees with recent studies showing that a recombinant TSEN34(R58W) mutant fails to hinder complex formation or activity but impacts the thermal stability of the complex^12^. Analysis of the thermal stability of additional PCH mutants combined with structural mapping of the mutants suggests that loss of TSEN complex stability may be an underlying cause of PCH^12^.

**Fig. 4.**
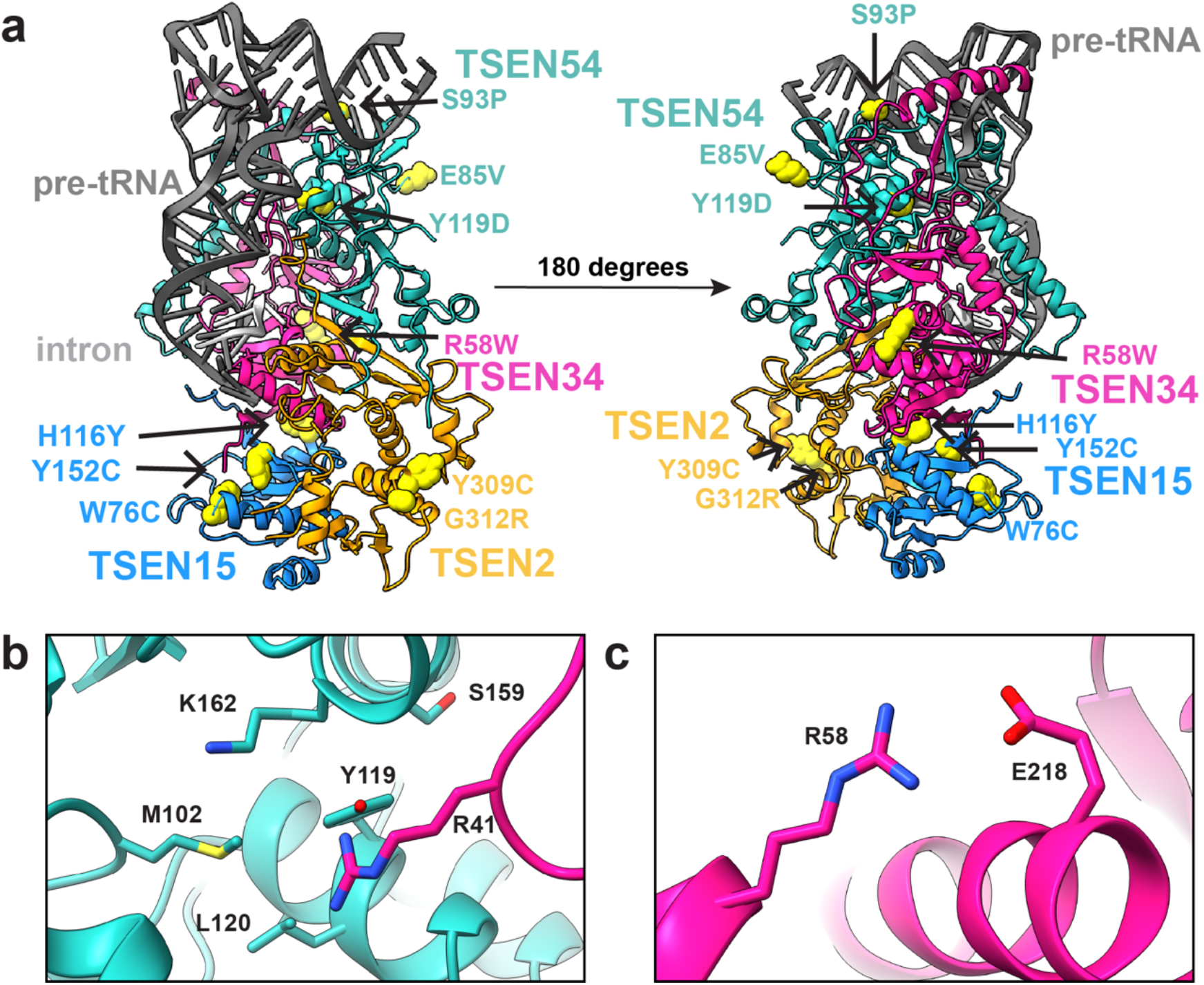
PCH Mutations mapped onto the TSEN complex structure. **a**. Previously reported PCH mutations are indicated with yellow spheres. **b**. TSEN54-Y119 is buried within the TSEN54 N-terminal domain and is in close proximity to TSEN34-R41 and several other TSEN54 (M102, L120, S159, K162) residues. **c**. TSEN34-R58 forms a salt bridge with TSEN34-E218.

## Summary

Our data provide significant insight into pre-tRNA recognition and processing by the human TSEN complex. The structure revealed an extensive tRNA-protein interface mediated by the TSEN54 subunit, which recognizes structural elements located in the acceptor stem, D-arm, and ASL of the pre-tRNA. Thus, instead of serving as a simple molecular ruler, TSEN54 works more like a set square that positions the tRNA along both sides of its L-shape. Beyond its capacity as a measuring device, TSEN54 also provides structural support for the two endonuclease subunits. Our structure also revealed many similarities and differences between archaeal and eukaryotic TSEN complexes. Both share a conserved core that facilitates recognition of the universal BHB motif and cleavage of the phosphodiester backbone. Beyond the conserved core, the structure revealed how additional regions of the eukaryotic TSEN subunits mediate pre-tRNA binding. The eukaryotic TSEN subunits also contain large unstructured regions which were not resolved in the structure and that could be important for supporting recruitment of other RNA substrates or mediating interactions with regulatory factors such as the CLP1 kinase. Finally, this study provides a mechanistic framework for understanding how mutations within the human TSEN complex impact its structure and cause PCH.

## Methods

### Preparation of wt-TSEN

The wt-TSEN complex was purified from *E*.*coli* as previously described using a multi-cistronic expression vector^17^. For BS3 crosslinking, the complex was purified over a Superdex-200 (Cytivia) with the following buffer: 50 mM HEPES pH 8.0, 200 mM NaCl, 5% glycerol, 5 mM MgCl_2_. For cryo-EM, the purified complex was subsequently incubated with tRNA-ARG-2’F (purchased HPLC purified from Horizon) and further purified over a Superdex-200 pre-equilibrated with 20 mM HEPES pH 8.0, 100 mM NaCl, 5 mM MgCl_2_, and 2% glycerol. Protein-tRNA fractions were pooled and concentrated and then diluted 1:1 with 20 mM HEPES pH 8.0, 100 mM NaCl, 5 mM MgCl_2_ buffer.

### Preparation of endoX-TSEN-CLP1 from HEK Cells

The endoX-TSEN•CLP1 complex was expressed and purified from large scale suspension cultures of HEK293 freestyle cells as previously described^17^, with the addition of 5 mM ATP to affinity resin washing steps and the following minor modifications: For cryo-EM, purification was performed using GFP nanobody resin^36^ and a C-terminal TEV-GFP-tag on CLP1. The TEV-tag was cleaved by overnight TEV digestion. The complex was then incubated with pre-tRNA-ARG, prepared as previously described^17^, and followed by Superdex-200 purification. Protein-tRNA complexes were prepared as described above for wt-TSEN. For BS3 crosslinking, CLP1 contained a C-terminal FLAG-Tag and the sample was purified using ANTI-FLAG M2 Affinity Gel (Sigma), eluted with Pierce 3X FLAG peptide (Thermo) and then further purified as described above for wt-TSEN.

### Protein-protein crosslinking

The TSEN•CLP1 (0.74 mg/mL) complex was crosslinked with 75 μM bis(sulfosuccinimidyl)suberate (BS3; Sigma). The wt-TSEN complex (1.6 mg/mL) was crosslinked with BS3 (75, 250, 500 μM). Crosslinking was quenched with Tris pH 8.0 to a final concentration of 30 mM on ice. Each crosslinked sample had a matching DMSO control. Samples were stored at -80 ºC until processing and analysis. Sample processing and interpretation was performed as previously described^37^.

### HEK cell immunoprecipitations and nuclease assays

Co-immunoprecipitation and pre-tRNA-ILE^TAT^-broccoli cleavage assays were conducted as previously described^17^ with minor differences in the overexpression constructs used. The constructs for the TSEN•CLP1 experiments were pLexM-CLP1–TEV-GFP, pCDNA3.1-TSEN15-Flag, pCDNA3.1-TSEN2-Flag, pCAG-Strep-Flag-TSEN34, pLexM-TSEN54-Flag. For the cation-π experiments, the constructs generated and cloned by Genscript into pCDNA3.1 vectors with a C-terminal (TSEN34) or N-terminal (TSEN2) Flag tag. The endonuclease triple mutants were cloned into pCAG-Strep-Flag-TSEN34 and pCDNA3.1-TSEN2-Flag. The non-bait TSEN proteins contained MYC tags.

### Cryo-EM grid preparation

Prior to freezing, grids were glow discharged (30 s, 15 mA at 0.37 mBar; Pelco Easiglow) and vitrified (3 μL sample, 3 s blot time) using an automatic plunge freezer (Leica). EndoX-TSEN•CLP1 was frozen on R1.2/1.3 Cu 300 mesh grids (Quantifoil) and TSEN (WT) was frozen on R2.4 Cu 300 mesh grids (Quantifoil).

### Cryo-EM Data Collection

TSEN micrographs were collected using SerialEM on a Talos Arctica electron microscope at 200 keV with a Gatan K2 Summit detector and a Titan Krios at 300 keV with a K3 Bioquantum detector. Beam-induced motion and drift were corrected using MotionCor2^38^ using Scipion3 software.

### Cryo-EM Image Processing

Image processing was conducted using CryoSPARC^39^. The CTF parameters of the dose-weighted images were calculated using CTFFIND4^40^. The complex 2D projections were initially selected using Blob Picker software and extracted with box size of 400. Picked particles were cleaned using a series of 2D Classifications and then a subset of cleaned particles were used for training template for Topaz Extraction which uses machine-learning. The Topaz picked particles were extracted using a box size of 320, with a down sampling factor of 2. Particles from different datasets of the same sample were then combined and further processed with additional iterations and selections of 2D Classifications. Ab-initio Model Reconstruction was then used to generate initial 3D models, which were used to further separate particles at the 3D level by Heterogeneous Refinements. The cleaned particles were then re-extracted to remove the down sampling factor, prior to final refinement steps. The 3D maps for both endoX-TSEN and wt-TSEN were further refined using unsymmetrized Homogeneous Refinement, Global CTF Refinement, Non-Uniform Refinement (endoX-TSEN only) and tight-masked Local Refinements.

### Model Building and Refinement

A combination of deposited PDB structures and alpha-fold models were used to construct the initial model of the TSEN complex and pre-tRNA. An unmodified tRNA-phe (PDB ID: 3L0U) was used as the starting point for building the pre-tRNA. The crystal structure of TSEN15-TSEN34 was used as the starting point for building TSEN15 (PDB ID: 6Z9U) and Alpha-fold models^24,25^ were used for TSEN34, TSEN54, and TSEN2. The starting models were individually docked into the reconstruction using UCSF Chimera^41^, followed by manual model building in Coot^42^. The structure was then refined through iterative rounds of manual model building and real-space refinement in Phenix^43^. The structure was validated using MolProbity^44^.

### Molecular Dynamics

Three different complexes were used in MD simulations; Complex (I) – TSEN34 (residues 1-103:181-309)/TSEN15 (residues 39-161)/TSEN2 (residues 39-71:295-464)/TSEN54 (residues 8-173:429-523) with the RNA present in the CryoEM structure (PDBID: XXXX), Complex (II) – all proteins from Complex (I) without the RNA, Complex (III) – all but TSEN15 from Complex (I). In the initial model of the RNA bound TSEN complex, only the missing segment of residues from 372 to 376 of TSEN2 was modeled in using the program, Modeller^45^. Protonation states of the histidine residues were selected using the web-based program, Molprobity^44^. All missing protons were introduced by using the leap module of Amber.18^46^. The complexes were solvated in a box of TIP3P water with the box boundary extending to 20 Å from the nearest protein atom (resulting in a total of 205708, 196569, and 204729 atoms in the rectangular simulation box of Complex (I), Complex (II), and Complex (III), respectively). Complex (I) contained 83 Na^+^ counter ions for neutralizing the charges while Complex (II) and Complex (III) had 2 and 74 Na+ ions, respectively. There were additional 116 Na^+^ and 116 Cl^-^ ions added to the solution of Complex (I) providing the 100 mM effective salt concentration approximating that at the physiological conditions whereas Complex (II) contained additional 112 Na+ and 112 Cl-ions and Complex (III) had 116 Na+ and 116 Cl-ions. Prior to equilibration, the solvated complexes were sequentially subjected to (i) 500 ps of dynamics of water and all ions with fixed (or frozen) proteins and RNA (if present), (ii) 5000 steps of energy minimization of all atoms (the first 2000 steps with the steepest decent method followed by the conjugate gradient method for the rest), (III) an initial constant temperature (300 K) - constant pressure (at 1 atm with the isotropic position scaling) dynamics at fixed protein to assure a reasonable starting density (∼1 ns) while keeping the protein positions under constraints with a 10 kcal/mol force constant for 50 ps, (iv) a conjugate-gradient minimization for 1000 steps, (v) step-wise heating MD at constant volume (from 0 to 300K in 3ns), and 6) constant volume simulation for 10 ns with a constraint force constant of 10 kcal/mol applied only on backbone heavy atoms. After releasing all constraining forces within the first 10 ns of the next 40 ns equilibration period, the constant-temperature (300 K) constant-volume (NVT) production runs were performed for a 1.0 µs. Sampling was increased by performing 5 independent 1.0 µs NVT molecular dynamics simulations for each of the three complexes, resulting in a total of 15 microsecond simulations. The constant temperature was maintained using the Langevin dynamics with the collision frequency of 0.5 ps^-1^. All trajectories were calculated using the PMEMD module of Amber.18 with 1 fs time step. Long-range coulombic interactions were handled using the PME method with the cut-off of 10 Å for the direct interactions. The amino acid parameters were selected from the FF14SB force field of Amber.18. RNA was represented by the FF99bsc0-chiOL3 force field. Root mean square deviations, root mean square fluctuations, dynamic cross correlation matrices, and various types of relevant analysis were performed using the CPPTRAJ module of Amber.18 in conjunction with some in-house programs. At the salt concentration of 100 mM with the standard parameters, the MM/GBSA module was used to estimate residue-residue interaction energies for all protein residues. A total of 5000 configurations was selected at each nanosecond from the five 1.0 µs MD trajectories for each of the complex studied.

## Data availability

Cryo-EM maps for wt-TSEN and endoX-TSEN have been deposited in the EMDB under the accession codes EMD-##### and EMD-26856 respectively. The atomic model for endoX-TSEN has been deposited in the PDB under the accession code PDBID-7UXA.

## Acknowledgements

We thank Drs. Percy Tumbale and Joseph Rodriguez for their critical reading of this manuscript. We would like to thank Dr. Rick Huang and Allison Zeher for help with cryo-EM data collection. This work utilized the Krios at the NCI/NICE Cryo-EM Facility. This work was supported by the US National Institute of Health Intramural Research Program; US National Institute of Environmental Health Sciences (ZIA ES103247 to R.E.S., 1ZIC ES102488 to J.G.W., 1ZIC ES103206 to L.J.D., 1ZIC ES102487 to R.M.P., 1ZI ES043010 to L.P., and 1ZIC ES103326 to M.J.B.). This work was also supported by the US National Institute of Health Extramural Research Program; US National Institute of General Medical Sciences (R35-GM136435, to A.G.M. and 1K99-GM143534 to C.K.H).

## Author Contributions

C.K.H., A.G.M., and R.E.S conceived and designed the study. C.K.H. and Z.D.S. prepared the recombinant *E. coli* TSEN sample and Z.D.S. performed relevant grid preparation. Z.D.S. also performed purification and crosslinking for mass spectrometry. J.G.W. and L.J.D. processed and analyzed mass spectrometry data. R.P. performed the HEK cell culture and transfections and supplied the anti-GFP resin. CKH prepared the endoX-TSEN-CLP1 complex sample, perform IP and nuclease assays, and generated tRNA-ARG. C.K.H. and K.J.B prepared endoX-TSEN-CLP1-tRNA-Arg grids and performed Cryo-EM screening. K.J.B. collected and pre-processed cryo-EM data. C.K.H. and K.J.B. performed all Cryo-EM analysis and structure determination. M.J.B. provided critical support in the design and processing of cryo-EM experiments. L.M.P performed all MD simulations. R.E.S, C.K.H., and J.M.K. performed structural refinement. C.K.H., R.E.S., and K.J.B. prepared the figures. C.K.H. and R.E.S. wrote and revised the manuscript.

## Competing interests

The authors declare no conflict of interest.

## Notes

### Competing Interest Statement

The authors have declared no competing interest.

## References

1 Hopper, A. K. & Nostramo, R. T. tRNA Processing and Subcellular Trafficking Proteins Multitask in Pathways for Other RNAs. Front Genet 10, 96, doi:10.3389/fgene.2019.00096 (2019).

2 Schmidt, C. A. & Matera, A. G. tRNA introns: Presence, processing, and purpose. Wiley Interdiscip Rev RNA 11, e1583, doi:10.1002/wrna.1583 (2020).

3 Gogakos, T. et al. Characterizing Expression and Processing of Precursor and Mature Human tRNAs by Hydro-tRNAseq and PAR-CLIP. Cell Rep 20, 1463–1475, doi:10.1016/j.celrep.2017.07.029 (2017).

4 Chan, P. P. & Lowe, T. M. GtRNAdb 2.0: an expanded database of transfer RNA genes identified in complete and draft genomes. Nucleic Acids Res 44, D184–189, doi:10.1093/nar/gkv1309 (2016).

5 Hayne, C. K., Lewis, T. A. & Stanley, R. E. Recent insights into the structure, function, and regulation of the eukaryotic transfer RNA splicing endonuclease complex. Wiley Interdiscip Rev RNA, e1717, doi:10.1002/wrna.1717 (2022).

6 Trotta, C. R. et al. The yeast tRNA splicing endonuclease: a tetrameric enzyme with two active site subunits homologous to the archaeal tRNA endonucleases. Cell 89, 849–858 (1997).

7 Paushkin, S. V., Patel, M., Furia, B. S., Peltz, S. W. & Trotta, C. R. Identification of a human endonuclease complex reveals a link between tRNA splicing and pre-mRNA 3’ end formation. Cell 117, 311–321, doi:10.1016/s0092-8674(04)00342-3 (2004).

8 Xue, S., Calvin, K. & Li, H. RNA recognition and cleavage by a splicing endonuclease. Science 312, 906–910, doi:10.1126/science.1126629 (2006).

9 Reyes, V. M. & Abelson, J. Substrate recognition and splice site determination in yeast tRNA splicing. Cell 55, 719–730, doi:10.1016/0092-8674(88)90230-9 (1988).

10 Greer, C. L., Soll, D. & Willis, I. Substrate recognition and identification of splice sites by the tRNA-splicing endonuclease and ligase from Saccharomyces cerevisiae. Mol Cell Biol 7, 76–84, doi:10.1128/mcb.7.1.76-84.1987 (1987).

11 Song, J. & Markley, J. L. Three-dimensional structure determined for a subunit of human tRNA splicing endonuclease (Sen15) reveals a novel dimeric fold. J Mol Biol 366, 155–164, doi:10.1016/j.jmb.2006.11.024 (2007).

12 Sekulovski, S. et al. Assembly defects of human tRNA splicing endonuclease contribute to impaired pre-tRNA processing in pontocerebellar hypoplasia. Nat Commun 12, 5610, doi:10.1038/s41467-021-25870-3 (2021).

13 Hirata, A. Recent Insights Into the Structure, Function, and Evolution of the RNA-Splicing Endonucleases. Front Genet 10, 103, doi:10.3389/fgene.2019.00103 (2019).

14 Yoshihisa, T. Handling tRNA introns, archaeal way and eukaryotic way. Front Genet 5, 213, doi:10.3389/fgene.2014.00213 (2014).

15 Calvin, K. & Li, H. RNA-splicing endonuclease structure and function. Cell Mol Life Sci 65, 1176–1185, doi:10.1007/s00018-008-7393-y (2008).

16 Weitzer, S., Hanada, T., Penninger, J. M. & Martinez, J. CLP1 as a novel player in linking tRNA splicing to neurodegenerative disorders. Wiley Interdiscip Rev RNA 6, 47–63, doi:10.1002/wrna.1255 (2015).

17 Hayne, C. K., Schmidt, C. A., Haque, M. I., Matera, A. G. & Stanley, R. E. Reconstitution of the human tRNA splicing endonuclease complex: insight into the regulation of pre-tRNA cleavage. Nucleic Acids Res 48, 7609–7622, doi:10.1093/nar/gkaa438 (2020).

18 Pacheva, I. H. et al. TSEN54 Gene-Related Pontocerebellar Hypoplasia Type 2 Could Mimic Dyskinetic Cerebral Palsy with Severe Psychomotor Retardation. Front Pediatr 6, 1, doi:10.3389/fped.2018.00001 (2018).

19 Schaffer, A. E. et al. CLP1 founder mutation links tRNA splicing and maturation to cerebellar development and neurodegeneration. Cell 157, 651–663, doi:10.1016/j.cell.2014.03.049 (2014).

20 Qian, Y. et al. A familial lateonset hereditary ataxia mimicking pontocerebellar hypoplasia caused by a novel TSEN54 mutation. Mol Med Rep 10, 1423–1425, doi:10.3892/mmr.2014.2342 (2014).

21 Karaca, E. et al. Human CLP1 mutations alter tRNA biogenesis, affecting both peripheral and central nervous system function. Cell 157, 636–650, doi:10.1016/j.cell.2014.02.058 (2014).

22 Laugwitz, L. et al. Pontocerebellar hypoplasia type 11: Does the genetic defect determine timing of cerebellar pathology? Eur J Med Genet 63, 103938, doi:10.1016/j.ejmg.2020.103938 (2020).

23 Forconi, M. et al. 2’-Fluoro substituents can mimic native 2’-hydroxyls within structured RNA. Chem Biol 18, 949–954, doi:10.1016/j.chembiol.2011.07.014 (2011).

24 Tunyasuvunakool, K. et al. Highly accurate protein structure prediction for the human proteome. Nature 596, 590–596, doi:10.1038/s41586-021-03828-1 (2021).

25 Jumper, J. et al. Highly accurate protein structure prediction with AlphaFold. Nature 596, 583–589, doi:10.1038/s41586-021-03819-2 (2021).

26 Li, H., Trotta, C. R. & Abelson, J. Crystal structure and evolution of a transfer RNA splicing enzyme. Science 280, 279–284, doi:10.1126/science.280.5361.279 (1998).

27 Abelson, J., Trotta, C. R. & Li, H. tRNA splicing. J Biol Chem 273, 12685–12688, doi:10.1074/jbc.273.21.12685 (1998).

28 Schmidt, C. A., Giusto, J. D., Bao, A., Hopper, A. K. & Matera, A. G. Molecular determinants of metazoan tricRNA biogenesis. Nucleic Acids Res 47, 6452–6465, doi:10.1093/nar/gkz311 (2019).

29 Tocchini-Valentini, G. D., Fruscoloni, P. & Tocchini-Valentini, G. P. Evolution of introns in the archaeal world. Proc Natl Acad Sci U S A 108, 4782–4787, doi:10.1073/pnas.1100862108 (2011).

30 Mattoccia, E., Baldi, I. M., Gandini-Attardi, D., Ciafre, S. & Tocchini-Valentini, G. P. Site selection by the tRNA splicing endonuclease of Xenopus laevis. Cell 55, 731–738, doi:10.1016/0092-8674(88)90231-0 (1988).

31 Fabbri, S. et al. Conservation of substrate recognition mechanisms by tRNA splicing endonucleases. Science (New York, N.Y.) 280, 284–286 (1998).

32 Fruscoloni, P., Baldi, M. I. & Tocchini-Valentini, G. P. Cleavage of non-tRNA substrates by eukaryal tRNA splicing endonucleases. EMBO Rep 2, 217–221, doi:10.1093/embo-reports/kve040 (2001).

33 Di Nicola Negri, E. et al. The eucaryal tRNA splicing endonuclease recognizes a tripartite set of RNA elements. Cell 89, 859–866, doi:10.1016/s0092-8674(00)80271-8 (1997).

34 Trotta, C. R., Paushkin, S. V., Patel, M., Li, H. & Peltz, S. W. Cleavage of pre-tRNAs by the splicing endonuclease requires a composite active site. Nature 441, 375–377, doi:10.1038/nature04741 (2006).

35 Baldi, M. I., Mattoccia, E., Bufardeci, E., Fabbri, S. & Tocchini-Valentini, G. P. Participation of the intron in the reaction catalyzed by the Xenopus tRNA splicing endonuclease. Science 255, 1404–1408, doi:10.1126/science.1542788 (1992).

36 Schellenberg, M. J., Petrovich, R. M., Malone, C. C. & Williams, R. S. Selectable high-yield recombinant protein production in human cells using a GFP/YFP nanobody affinity support. Protein Sci 27, 1083–1092, doi:10.1002/pro.3409 (2018).

37 Pillon, M. C. et al. Cryo-EM reveals active site coordination within a multienzyme pre-rRNA processing complex. Nature structural & molecular biology 26, 830–839, doi:10.1038/s41594-019-0289-8 (2019).

38 Zheng, S. Q. et al. MotionCor2: anisotropic correction of beam-induced motion for improved cryo-electron microscopy. Nat Methods 14, 331–332, doi:10.1038/nmeth.4193 (2017).

39 Punjani, A., Rubinstein, J. L., Fleet, D. J. & Brubaker, M. A. cryoSPARC: algorithms for rapid unsupervised cryo-EM structure determination. Nat Methods 14, 290–296, doi:10.1038/nmeth.4169 (2017).

40 Rohou, A. & Grigorieff, N. CTFFIND4: Fast and accurate defocus estimation from electron micrographs. J Struct Biol 192, 216–221, doi:10.1016/j.jsb.2015.08.008 (2015).

41 Pettersen, E. F. et al. UCSF Chimera--a visualization system for exploratory research and analysis. J Comput Chem 25, 1605–1612, doi:10.1002/jcc.20084 (2004).

42 Emsley, P., Lohkamp, B., Scott, W. G. & Cowtan, K. Features and development of Coot. Acta Crystallogr D Biol Crystallogr 66, 486–501, doi:10.1107/S0907444910007493 (2010).

43 Adams, P. D. et al. PHENIX: a comprehensive Python-based system for macromolecular structure solution. Acta Crystallogr D Biol Crystallogr 66, 213–221, doi:10.1107/S0907444909052925 (2010).

44 Williams, C. J. et al. MolProbity: More and better reference data for improved all-atom structure validation. Protein Sci 27, 293–315, doi:10.1002/pro.3330 (2018).

45 Webb, B. & Sali, A. Comparative Protein Structure Modeling Using MODELLER. Curr Protoc Bioinformatics 54, 5 6 1–5 6 37, doi:10.1002/cpbi.3 (2016).

46 D.A. Case, I. Y. B.-S., S.R. Brozell, D.S. Cerutti, T.E. Cheatham, III, V.W.D. Cruzeiro, T.A. Darden, R.E. Duke, D. Ghoreishi, M.K. Gilson, H. Gohlke, A.W. Goetz, D. Greene, R Harris, N. Homeyer, Y. Huang, S. Izadi, A. Kovalenko, T. Kurtzman, T.S. Lee, S. LeGrand, P. Li, C. Lin, J. Liu, T. Luchko, R. Luo, D.J. Mermelstein, K.M. Merz, Y. Miao, G. Monard, C. Nguyen, H. Nguyen, I. Omelyan, A. Onufriev, F. Pan, R. Qi, D.R. Roe, A. Roitberg, C. Sagui, S. Schott-Verdugo, J. Shen, C.L. Simmerling, J. Smith, R. Salomon-Ferrer, J. Swails, R.C. Walker, J. Wang, H. Wei, R.M. Wolf, X. Wu, L. Xiao, D.M. York and P.A. Kollman AMBER 2018. University of California, San Francisco. (2018).

