## Supplemental for "Structural Basis for pre-tRNA Recognition and Processing by the Human tRNA Splicing Endonuclease Complex"

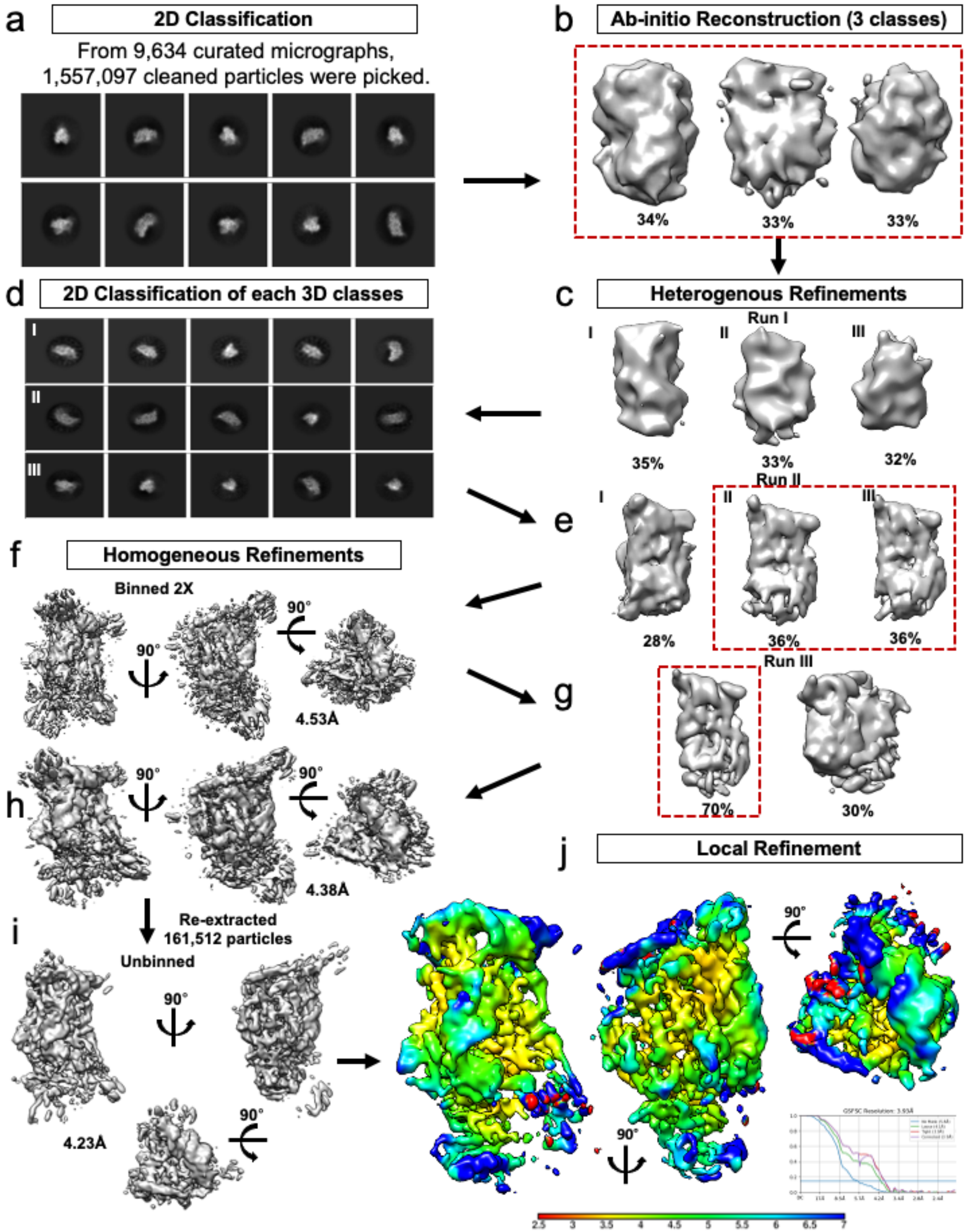


**Extended Data Fig. 1 | WT-TSEN cryo-EM processing workflow**. **a**. 1,557,097 particles were picked from 9,634 curated micrographs, 10 representative 2D classes are shown. Particles contained in good classes were used to generate **b**. ab-initio reconstructions, three of which were selected and further refined using **c**. heterogenous refinement. The particles contained in the best three classes were further filtered using **d**. 2D classification and **e**. heterogeneous refinement prior to two classes being further refined using **f.** homogenous refinement, with 2x binned particles. **g.** A final round of heterogenous refinement resulted in one 3D class which was **h.** refined to an estimated 4.38 Å using homogenous refinement. **i**. Refinement continued with another round of homogeneous refinement resulting in a resolution of 4. Å following the unbinning of the final 161,512 particles with an estimated resolution of 4.2 Å. **j**. A local refinement resulted in a final map with an estimated resolution of 3.9 Å.

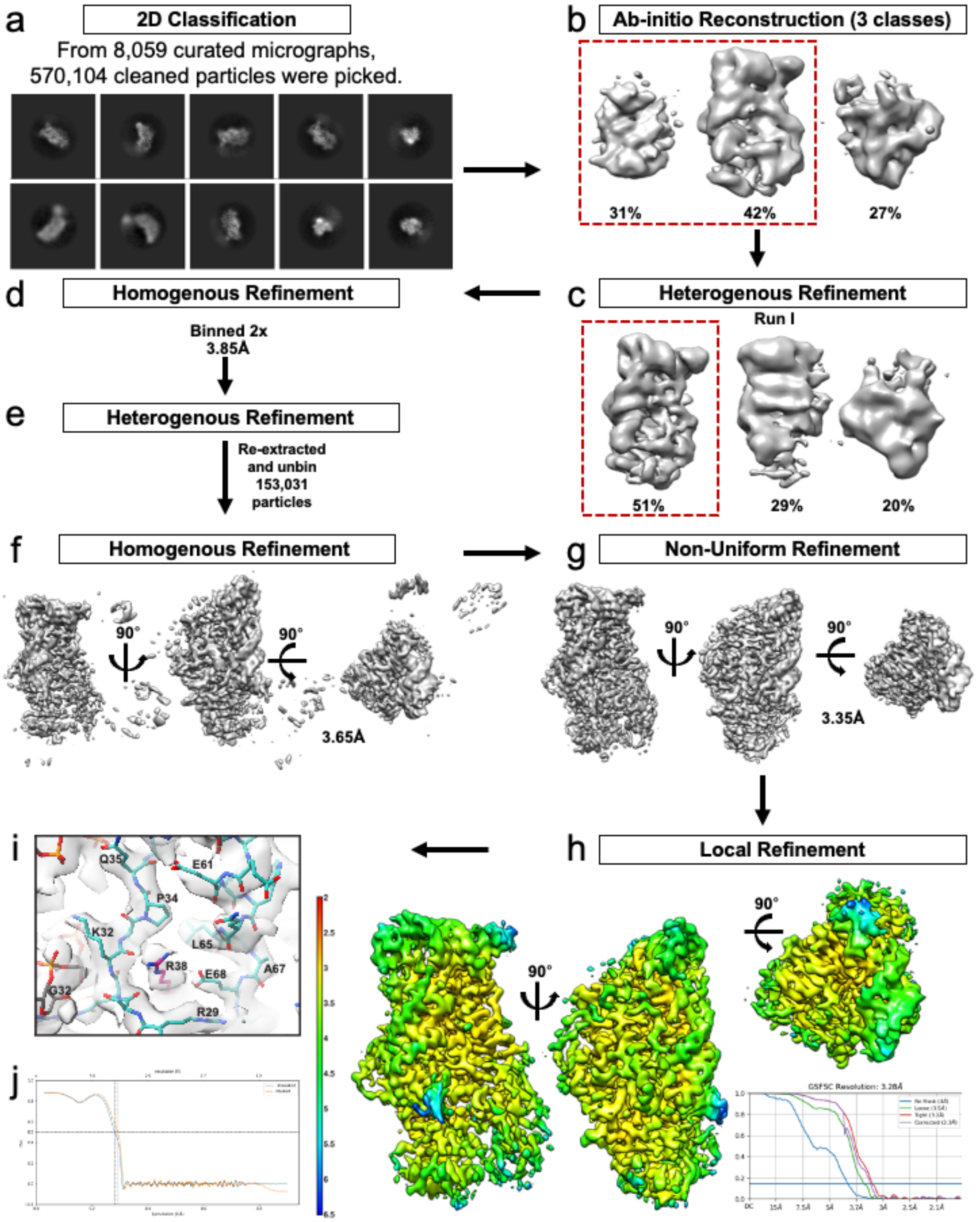


**Extended Data Fig. 2 | EndoX-TSEN structure cryo-EM processing workflow**.

**a**. 570,104 particles were picked from 8059 curated micrographs and used for 2D classification. Particles from the good classes were used to generate **b**. ab-initio reconstructions, two of which were selected and further refined using **c**. heterogenous refinement. **d**. The particles contained in the best class were binned and refined with homogenous refinement. **e**. The particles were then reextracted and unbinned and used for another round of **f.** homogenous refinement followed by **g.** non-uniform refinement and **h**. local refinement resulting in a final map with an estimated resolution of 3.28 Å. **i.** Example density for the complex, showing a region of TSEN54's N-terminus (39-35, 61-67) and TSEN34-R38 **j.** FSC of the model to map

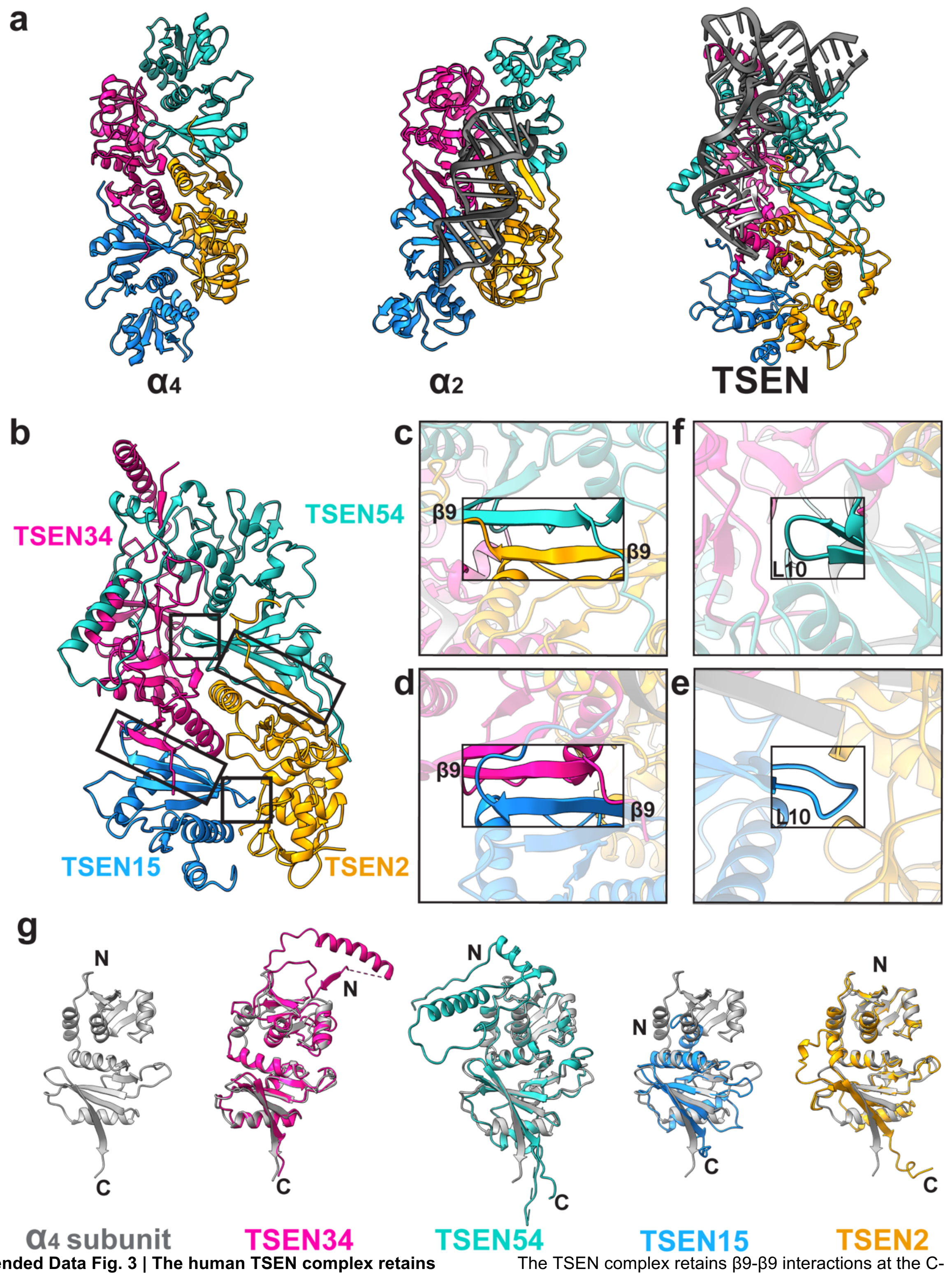


**Extended Data Fig. 3 | The human TSEN complex retains the core architecture from the archaeal complex. a**. Structures of a homotetrameric (α4, PDBID: 1A79), homodimeric BHB bound (α2, PDBID:2GJW), and the pre-tRNA bound TSEN complex, colored to mimic the arrangement of the subunits within the TSEN complex (TSEN54 – teal, endonuclease TSEN2- orange, endonuclease TSEN34 –pink, TSEN15 – medium blue). **b**. The TSEN complex retains β9-β9 interactions at the C-terminal domain of endonuclease/structural protein interfaces: **c**. TSEN2:TSEN54 and **d**. TSEN34:TSEN15. The L10-loop of the structural proteins **e**. TSEN54 and **f**. TSEN15 link the β9-β9 heterodimers together. **g**. A single archaeal homotetramer α subunit (PDBID: 1A79, grey) superimposed with each individual TSEN subunit.

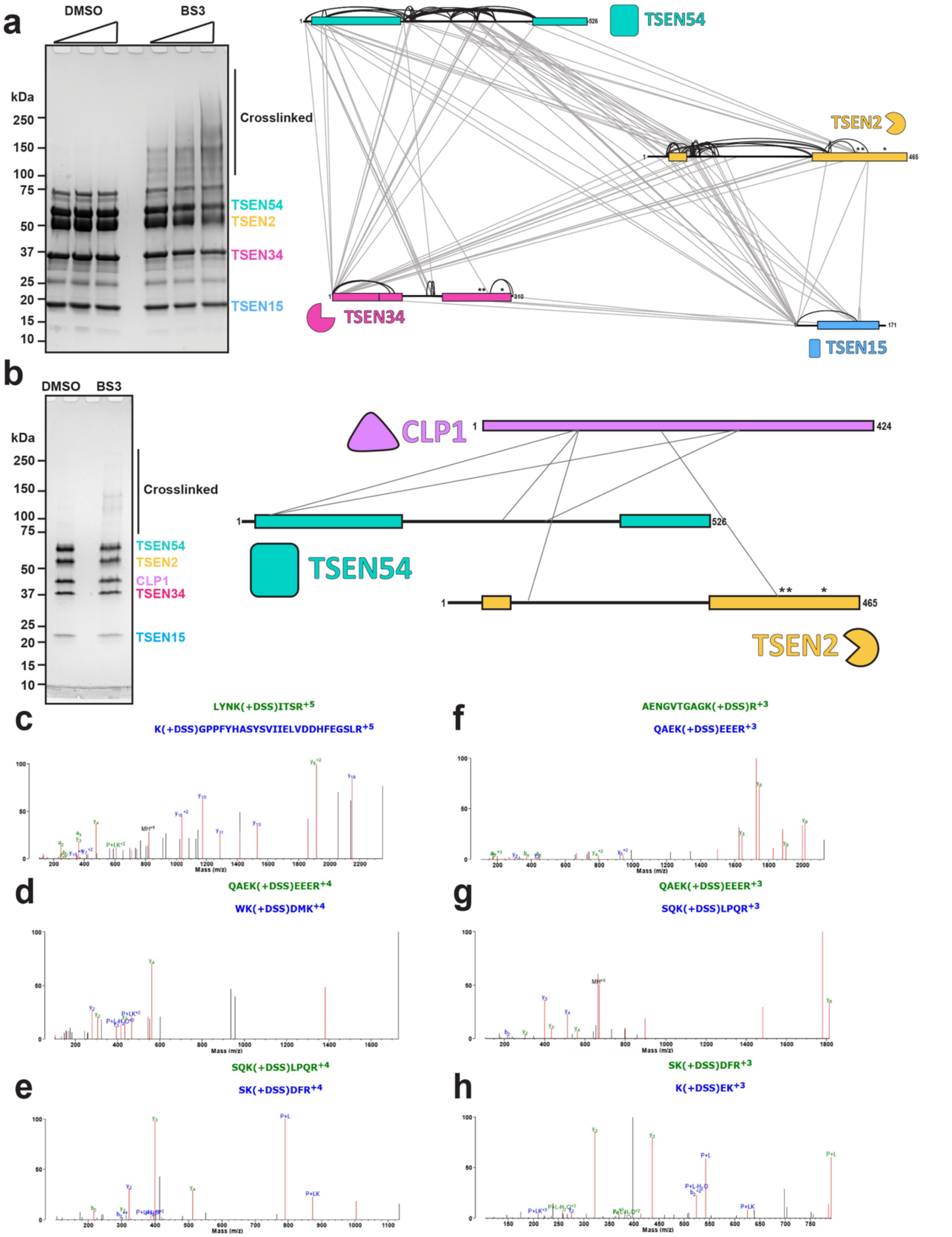


**Extended Data Fig. 4 | BS3 crosslinking of TSEN complexes.**

SDS-Page gels showing the DMSO control and crosslinked samples for **a.** wt-TSEN complex and **b**. TSEN Complex and CLP1. **c-h.** MSMS spectra of CLP1 peptides crosslinked to other members of the complex. Panels c (m/z 822.4292) and d (466.4795 m/z) show MSMS spectra of peptides arising from CLP1 and TSEN2. Panels e (m/z 412.2311), f (m/z 739.0369), g (m/z 671.3516), and h (m/z 398.5568) show MSMS spectra arising from crosslinks from CLP1 and TSEN54. Fragment ions that arise from green colored peptides are shown in green, fragments arising from blue colored peptides are shown in blue, and unassigned ions shown in black.

**a**

**Root mean square deviations**

| Complex (I) | run 1 | run 2 | run 3 | run 4 | run 5 | average |
|---|---|---|---|---|---|---|
| TSEN34 | 2.39(0.26) | 2.41(0.28) | 3.76(0.73) | 3.49(0.43) | 2.39(0.38) | 2.89(0.61) |
| TSEN15 | 1.22(0.12) | 1.43(0.21) | 1.20(0.13) | 1.62(0.27) | 1.32(0.15) | 1.36(0.16) |
| TSEN2 | 2.18(0.25) | 1.88(0.17) | 1.79(0.13) | 2.28(0.17) | 1.86(0.21) | 2.00(0.20) |
| TSEN54 | 1.81(0.17) | 2.13(0.22) | 1.56(0.14) | 1.66(0.15) | 2.29(0.26) | 1.89(0.28) |
| Total | 2.42(0.22) | 2.57(0.25) | 2.94(0.38) | 3.29(0.34) | 2.58(0.38) | 2.76(0.32) |

| Complex (II) | run 1 | run 2 | run 3 | run 4 | run 5 | average |
|---|---|---|---|---|---|---|
| TSEN34 | 2.49(0.30) | 2.90(0.59) | 4.18(0.62) | 2.18(0.36) | 2.28(0.23) | 2.81(0.73) |
| TSEN15 | 1.33(0.19) | 1.31(0.15) | 1.23(0.11) | 1.24(0.14) | 1.51(0.28) | 1.32(0.10) |
| TSEN2 | 2.33(0.35) | 1.94(0.18) | 2.06(0.34) | 2.08(0.21) | 2.13(0.16) | 2.11(0.13) |
| TSEN54 | 2.04(0.20) | 1.82(0.16) | 2.00(0.22) | 1.90(0.19) | 2.13(0.20) | 1.98(0.11) |
| Total | 2.84(0.31) | 2.77(0.33) | 3.86(0.51) | 2.39(0.24) | 2.48(0.17) | 2.87(0.52) |

| Complex (III) | run 1 | run 2 | run 3 | run 4 | run 5 | average |
|---|---|---|---|---|---|---|
| TSEN34 | 3.13(0.44) | 4.25(0.47) | 1.82(0.19) | 2.17(0.34) | 2.12(0.21) | 2.70(0.89) |
| TSEN2 | 2.19(0.22) | 2.04(0.16) | 2.20(0.19) | 1.96(0.29) | 1.69(0.15) | 2.02(0.19) |
| TSEN54 | 1.70(0.16) | 1.42(0.09) | 1.45(0.13) | 1.46(0.12) | 1.62(0.14) | 1.53(0.11) |
| Total | 2.80(0.29) | 3.13(0.28) | 2.28(0.24) | 2.46(0.31) | 2.09(0.19) | 2.55(0.37) |

**b**

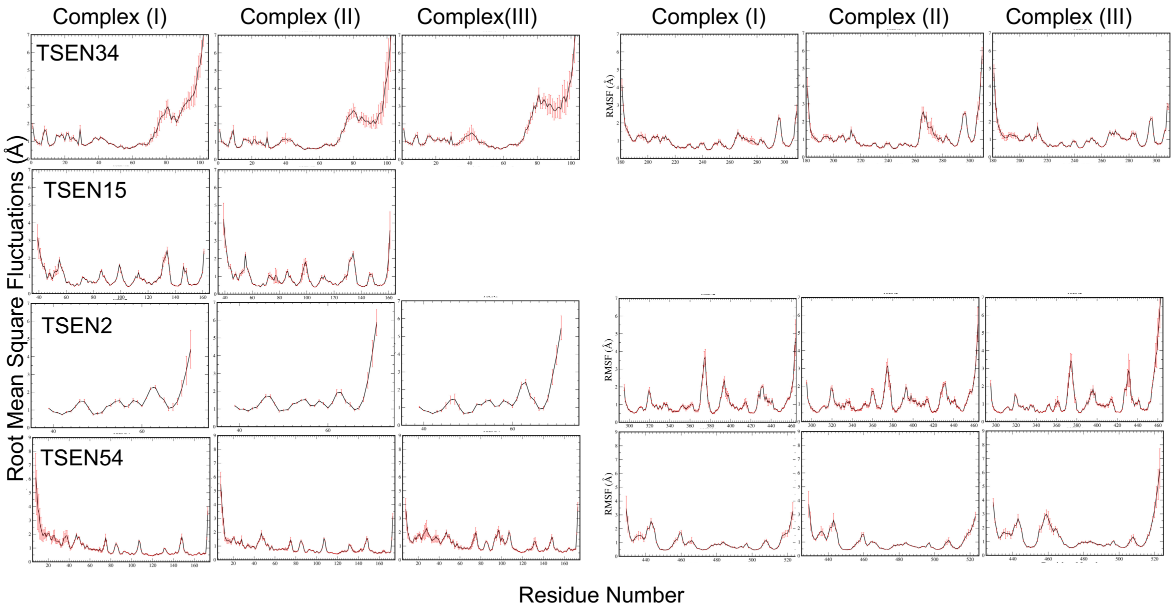


**c**

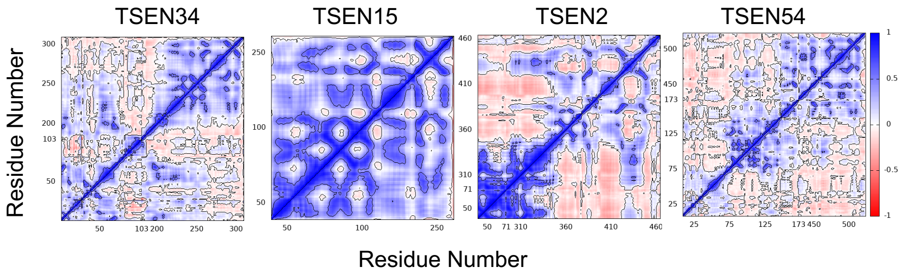


**Extended Data Fig. 5 | Molecular dynamics simulations account for stable complexes during dynamics**. **a.** Root mean square deviations (RMSD) of individual proteins and the complex averaged over each run and averaged over the five runs. TSEN34 has the largest deviations while TSEN15 shows smallest deviations irrespective of the RNA binding. Standard deviations are shown in parenthesis. The reference (Complex I) was the starting CryoEM configuration. Complex (II) is without tRNA and Complex (III) is without TSEN15. **b.** Root mean square fluctuations of individual residues averaged calculated during the microsecond dynamics and averaged over all runs. Standard deviations are shown as error bars. **c.** Representative dynamic cross correlation matrices of the protein components from Complex (I).

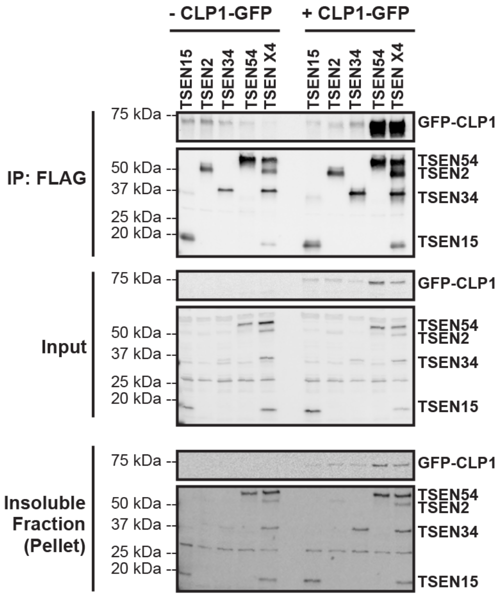


**Extended Data Fig. 6 | TSEN54 is the anchor that mediates the interaction of CLP1 with the TSEN complex.** Overexpression (in HEK cells) and immunoprecipitations of the individual TSEN proteins (lanes 1-4 and lanes 7-10) or full TSEN complex (lanes 5 and 11) in the absence (lanes 1-5) and presence (lanes 7-11) of CLP1 reveals strong association between CLP1 and TSEN54 (lane 10) as well as the full TSEN complex (lane 11). The experiments were conducted as previously reported[1] using CLP1-TEV-GFP.

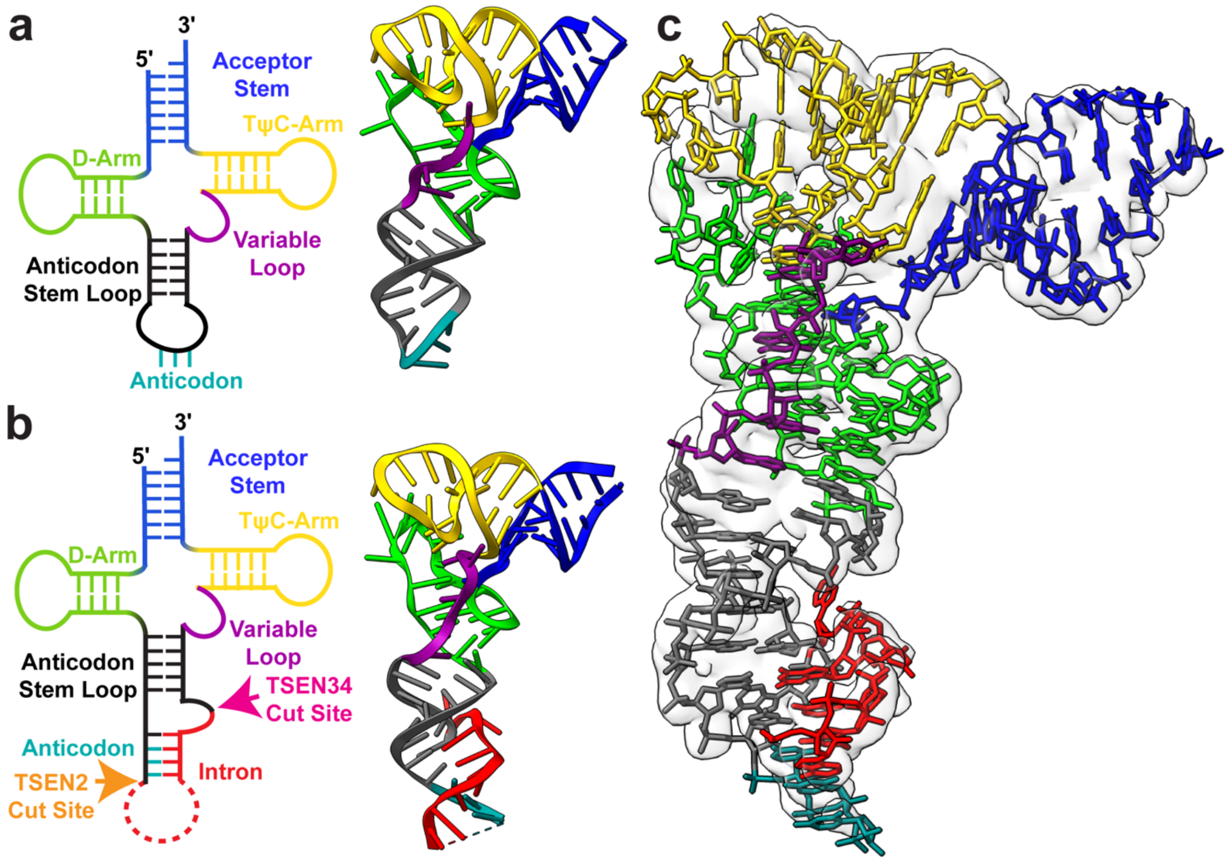


**Extended Data Fig. 7 | Structure of the pre-tRNA reveals a familiar architecture to mature tRNA.** 2D and 3D structures of **a.** unmodified mature tRNA-PHE (PDBID: 3L0U) and **b.** intron containing pre-tRNA-ARG structure, with the acceptor stem (blue), D-Arm (lime green), anticodon stem loop (dark grey), anticodon (teal), intron (red), variable loop (purple) and TψC-Arm (yellow) labeled. The cut sites for TSEN2 (orange) and TSEN34 (pink) are labelled on the pre-tRNA 2D structure. **c**. Electron density map around the pre-tRNA, with the pre-tRNA structure docked into the density map.

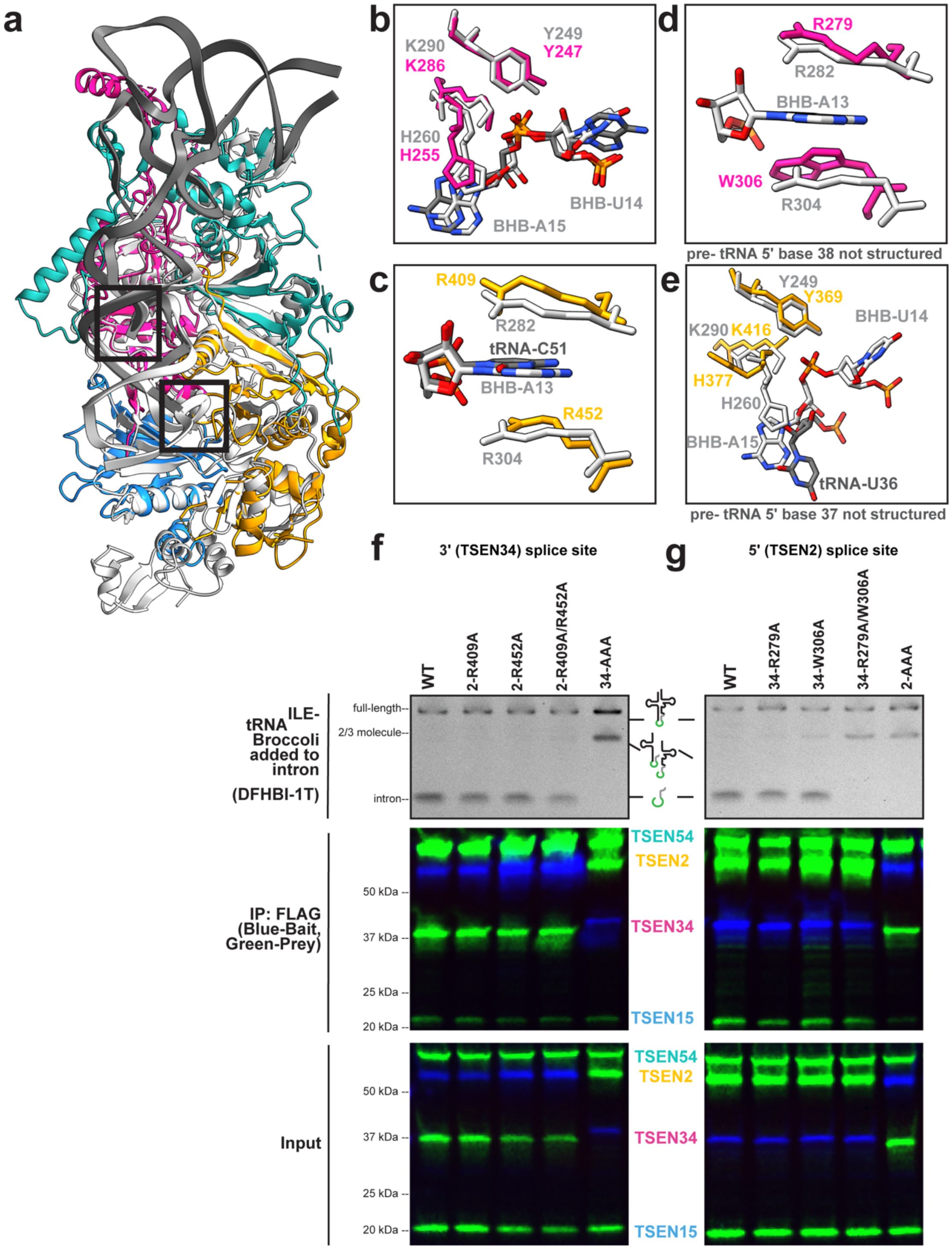


**Extended Data Fig. 8 | Structural comparison of the TSEN and EndA active site residues.**
**a**. Overlay of TSEN + tRNA (colored as before) and EndA + BHB (grey, PDBID:2GJW) showing overall similar RNA binding. **b**. Overlay of the active site residues for the 3' splice site and the **c**. cation-$\pi$ residues from TSEN2 shown. **d**. Overlay of the cation-$\pi$ residues from TSEN34 and the **e**. TSEN2 active site residues for the 5' splice site. **f.** tRNA cleavage assays of a broccoli RNA -aptamer containing pre-tRNA-ILE using samples immunoprecipitated via the mutants of the 3' splice site (pulled down by a FLAG tag on TSEN2, except for the 34 mutant) or **g.** 5' splice site (pulled down by a FLAG tag on TSEN34, except for TSEN2 active site mutant). TSEN proteins that weren't the pull-down target had a MYC-tag.

| | CLP1-TSEN(endoX)-<br>tRNA-Arg | | TSEN(WT)-<br>2’F-tRNA-Arg |
|---|---|---|---|
| **EMDB** | EMD-26856 | | |
| **PDB ID** | PDBID-7UXA | | N/A |
| **Data collection and processing** | | | |
| Microscope | Talos Arctica | Titan Krios | Titan Krios |
| Detector | Gatan K2 Summit | Gatan K3 Bioquantum | Gatan K3 Bioquantum |
| Magnification | 45000x | 81000x | 81000x |
| Voltage(kV) | 200 | 300 | 300 |
| Electron exposure (e−/Å2) | 54 | 60 | 60 |
| Defocus range (μm) | -0.8 to -1.8 | -1.2 to -2.2 | -1.2 to -2.2 |
| Pixel size (Å) | 0.93 | 0.53 | 0.53 |
| Symmetry imposed | C1 | | C1 |
| Number of micrographs | 8,059 | | 9,634 |
| Initial particle images (no.) | 570,104 | | 1,557,097 |
| Final particle images (no.) | 153,031 | | 161,512 |
| Map resolution (Å) (FSC = 0.143) | 3.28 | | 3.93 |
| Local Resolution Range (Å) | 3.31 | | 3.91 |
| FSC model (0/0.143/0.5) (Å) | 3.2/3.2/3.4 | | |
| **Refinement** | | | |
| Initial models used (PDB code) | 6Z9U, 3L0U | | |
| Map sharpening B factor ($Å^2$) | 121.9 | | |
| *Model composition* | | | |
| Nonhydrogen atoms | 8,120 | | |
| Protein residues | 820 | | |
| Nucleotides | 78 | | |
| Ligands | 2 (Mg) | | |
| *B factors (Å2)* | | | |
| Protein | 76.24 | | |
| Nucleotide | 81.90 | | |
| Ligand | 22.87 | | |
| *R.M.S. deviations* | | | |
| Bond lengths (Å) | 0.003 (0) | | |
| Bond angles (°) | 0.575 (0) | | |
| *Map-model CC* | | | |
| CC (mask) | 0.82 | | |
| CC (box) | 0.78 | | |
| CC (peaks) | 0.73 | | |
| CC (volume) | 0.81 | | |
| **Validation** | | | |
| Molprobity score | 1.78 | | |
| Clashscore | 7.47 | | |
| Poor rotamers (%) | 0.88 | | |
| *Ramachandran plot* | | | |
| Outliers (%) | 0.00 | | |
| Allowed (%) | 5.34 | | |
| Favored (%) | 94.66 | | |
| CaBLAM outliers (%) | 3.03 | | |

**Extended Data Table 1 |Cryo-EM data collection, refinement, and validation statistics.**

**a. Residue (nucleotide) interaction**

| RNA | |
|---|---|
| U4 | -3.09(0.15) |
| C5 | -9.06(0.31) |
| U6 | -12.28(0.47) |
| G7 | -10.18(0.35) |
| U8 | -4.21(0.26) |
| C11 | -18.37(0.45) |
| G12 | -15.87(0.43) |
| C13 | -20.53(0.57) |
| A14 | -20.41(0.70) |
| A15 | -10.23(0.65) |
| U16 | -24.43(1.30) |
| G17 | -38.47(1.40) |
| G22 | -11.51(0.78) |
| C23 | -16.94(0.85) |
| G24 | -12.01(0.80) |
| C25 | -18.24(0.72) |
| A26 | -12.00(0.71) |
| U27 | -3.76(0.71) |
| U33 | -7.52(0.63) |
| U34 | -16.58(0.90) |
| C35 | -16.96(0.91) |
| U36 | -13.28(0.91) |
| A44 | -4.22(0.46) |
| A45 | -4.51(0.46) |
| G48 | -5.91(0.35) |
| A49 | -13.10(0.80) |
| G50 | -18.68(1.14) |
| C51 | -36.47(1.18) |
| A52 | -31.82(0.96) |
| A53 | -45.97(0.99) |
| U54 | -16.88(0.77) |
| U55 | -15.80(0.95) |
| C56 | -12.89(0.80) |
| A57 | -3.54(0.61) |
| G83 | -6.84(0.53) |
| A84 | -25.96(0.80) |
| G85 | -33.86(1.12) |
| U86 | -9.22(0.35) |

| TSEN2 | |
|---|---|
| TYR377 | -3.11(0.35) |
| HIS378 | -4.32(0.35) |
| ARG410 | -21.27(0.72) |
| VAL413 | -5.52(0.22) |
| ASN414 | -10.72(0.48) |
| VAL415 | -4.69(0.21) |
| SER416 | -4.39(0.30) |
| LYS417 | -5.31(0.51) |
| ARG453 | -22.20(1.23) |
| SER457 | -6.20(0.34) |
| ARG458 | -18.55(1.40) |
| ARG460 | -21.47(0.65) |
| SER461 | -4.37(0.32) |

| TSEN54 | |
|---|---|
| ARG27 | -26.40(1.07) |
| ARG29 | -10.60(0.86) |
| GLN31 | -12.39(0.62) |
| LYS32 | -14.25(0.92) |
| GLN35 | -4.53(0.44) |
| ARG36 | -28.35(1.47) |
| LYS41 | -20.62(0.84) |
| ARG73 | -19.36(1.04) |
| PRO94 | -3.74(0.17) |
| GLY96 | -3.93(0.19) |
| LYS97 | -48.21(0.91) |
| TRP99 | -5.04(0.23) |
| GLN100 | -10.55(0.58) |
| VAL458 | -5.97(0.24) |
| ARG462 | -23.47(1.11) |
| LYS463 | -11.29(0.37) |
| ASN464 | -18.31(0.72) |

| TSEN34 | |
|---|---|
| ARG31 | -4.16(0.33) |
| ARG41 | -11.95(0.48) |
| GLN42 | -5.43(0.19) |
| SER44 | -3.01(0.25) |
| ARG45 | -9.06(0.82) |
| SER181 | -9.74(0.94) |
| LYS239 | -21.72(0.76) |
| HID255 | -12.74(0.32) |
| ARG279 | -11.12(0.99) |
| SER283 | -4.11(0.29) |
| VAL284 | -4.07(0.17) |
| ARG285 | -18.25(1.09) |
| LYS286 | -13.43(0.66) |

**b. RNA contact frequency**

| RNA | |
|---|---|
| U4 | 0.93(0.01) |
| C5 | 1.00(0.00) |
| U6 | 1.00(0.00) |
| G7 | 1.00(0.00) |
| U8 | 0.97(0.00) |
| G10 | 0.98(0.01) |
| C11 | 1.00(0.00) |
| G12 | 1.00(0.00) |
| C13 | 1.00(0.00) |
| A14 | 1.00(0.00) |
| A15 | 0.93(0.03) |
| U16 | 0.97(0.02) |
| G17 | 1.00(0.00) |
| G18 | 0.12(0.09) |
| A19 | 0.09(0.06) |
| G22 | 0.99(0.00) |
| C23 | 1.00(0.00) |
| G24 | 1.00(0.00) |
| C25 | 1.00(0.00) |
| A26 | 1.00(0.00) |
| U27 | 0.75(0.05) |
| C32 | 0.05(0.02) |
| U33 | 0.82(0.08) |
| U34 | 0.96(0.04) |
| C35 | 1.00(0.00) |
| U36 | 0.98(0.01) |
| A44 | 0.47(0.15) |
| A45 | 0.36(0.16) |
| U46 | 0.26(0.16) |
| A47 | 0.53(0.15) |
| G48 | 0.98(0.01) |
| A49 | 0.98(0.01) |
| G50 | 0.96(0.04) |
| C51 | 1.00(0.00) |
| A52 | 1.00(0.00) |
| A53 | 1.00(0.00) |
| U54 | 0.71(0.17) |
| U55 | 0.89(0.08) |
| C56 | 0.89(0.09) |
| A57 | 0.48(0.14) |
| C76 | 0.05(0.03) |
| G83 | 0.95(0.01) |
| A84 | 1.00(0.00) |
| G85 | 1.00(0.00) |
| U86 | 0.99(0.00) |
| C87 | 0.52(0.08) |

**C. Interaction strengths among different components in the complex**

| | Complex (I) | Complex (II) | Complex (III) |
|---|---|---|---|
| TSEN34-TSEN15 | -104.64(1.73) | -97.22(1.05) | --- |
| TSEN34-TSEN2 | -7.09(1.04) | -19.16(2.55) | -10.65(1.57) |
| TSEN34-TSEN54 | -224.16(3.48) | -246.32(4.19) | -224.78(3.18) |
| TSEN15-TSEN2 | -10.46(1.84) | -23.25(1.68) | --- |
| TSEN2-TSEN54 | -145.11(4.07) | -123.69(2.75) | -148.71(2.20) |
| Complex - RNA | -240.97(6.46) | --- | -240.65(5.70) |

**Extended Data Table 2: Energy contributions and contact frequencies from Molecular Dynamic simulations. a.** MMGBSA energy calculated from 5000 configurations collected from 5 MD runs. The standard errors are given in parenthesis. Only energies with magnitudes greater than 3 kcals/mol are shown. **b**. Contact frequencies of nucleotides with the TSEN complex. The frequencies were calculated from 5000 configurations and if any heavy atom of a nucleotide is within 3.2 A to any heavy atoms of the protein complex, then the nucleotide is selected to be in contact with the complex. Only 35 out of 80 nucleotides are in contact 75% or more of the time of the simulation. **c**. interaction energies among various components in each system, calculated using MMGBSA.
